# Characterization of Multicellular Niches Supporting Hematopoietic Stem Cells Within Distinct Zones

**DOI:** 10.1101/2024.06.28.601225

**Authors:** Ruochen Dong, Hua Li, Xi C He, Chen Wang, Anoja Perera, Seth Malloy, Jonathon Russell, Wenting Li, Kaitlyn Petentler, Xinjian Mao, Zhe Yang, Michael Epp, Kate Hall, Allison Scott, Mary C. McKinney, Shengping Huang, Sarah E Smith, Mark Hembree, Yongfu Wang, Zulin Yu, Jeffery S. Haug, Jay Unruh, Brian Slaughter, Xunlei Kang, Linheng Li

## Abstract

Previous studies of hematopoietic stem cells (HSCs) primarily focused on single cell-based niche models, yielding fruitful but conflicting findings^1–5^. Here we report our investigation on the fetal liver (FL) as the primary fetal hematopoietic site using spatial transcriptomics. Our study reveals two distinct niches: the portal-vessel (PV) niche and the sinusoidal niche. The PV niche, composing N-cadherin (N-cad)^Hi^Pdgfrα^+^ mesenchymal stromal cells (MSCs), endothelial cells (ECs), and N-cad^Lo^Albumin^+^ hepatoblasts, maintains quiescent and multipotential FL-HSCs. Conversely, the sinusoidal niche, comprising ECs, hepatoblasts and hepatocytes, as well as potential macrophages and megakaryocytes, supports proliferative FL-HSCs biased towards myeloid lineages. Unlike prior reports on the role of Cxcl12, with its depletion from vessel-associated stromal cells leading to 80% of HSCs’ reduction in the adult bone marrow (BM)^6,7^, depletion of *Cxcl12* via *Cdh2^CreERT^* (encoding N-cad) induces altered localization of HSCs from the PV to the sinusoidal niches, resulting in an increase of HSC number but with myeloid-bias. Similarly, we discovered that adult BM encompasses two niches within different zones, each composed of multi-cellular components: trabecular bone area (TBA, or metaphysis) supporting deep-quiescent HSCs, and central marrow (CM, or diaphysis) fostering heterogenous proliferative HSCs. This study transforms our understanding of niches by shifting from single cell-based to multicellular components within distinct zones, illuminating the intricate regulation of HSCs tailored to their different cycling states.

## Introduction

Hematopoietic stem cells (HSCs) possess the remarkable self-renew ability and multilineage potential, ensuring a lifelong supply of both HSCs and lineage cells^8^. Definitive HSCs emerge from the hemogenic endothelium in the aorta-gonad-mesonephros (AGM) region around E10.5-11.5 and subsequently migrate to the FL after E12.5 for expansion. The FL serves as the primary site for HSC maturation and expansion^9–12^. It is estimated that HSCs undergo a 38-fold expansion between E12 and E18 in the FL before migrating to the BM near birth to support lifelong hematopoiesis in adults^13–15^.

In the adult BM, it was proposed that HSCs reside within physically confined microenvironment or niche^16^, however the identification of the cell types that serve as niches in supporting HSCs within the BM has been a long journey over the past two decades^1–4,17^. The first HSC niche was found in the TBA or metaphysis as evidenced by the observations that increased TBA led to an increased HSC number^18,19^, thus forming the concept of the endosteal (inner bone surface) niche. Notably, in these studies, long-term (LT) quiescent HSCs were identified using two criteria: 1) LT BrdU labeling (for 10-days) followed by a 70-day chase to detect label-retaining cells (LRCs), and 2) HSC markers Lin^-^CD45^+^Sca1^+^Kit^+^ expression in a subset of LRCs. Within the increased TBA, more N-cad^+^ bone-lining cells correlates with more LRC^+^Lin^-^ CD45^+^Sca1^+^Kit^+^ HSCs^18,20^. It is important to acknowledge that employing different LRC procedures, including variations in labeling and chasing time, may result in the identification of different cell status or even cell types^21^. This is because each division of HSCs diminishes their labels such as BrdU or GFP and their self-renewal capacity, and it should be noted that not all LRCs necessarily correspond to HSCs^22,23^. Moreover, these LT quiescent HSCs were primarily found in the endosteal region (70-75%) and within the TBA, many of which were observed adjacent to N-cad^+^ bone-lining cells, although the exact nature of the latter was not fully understood at the time^18,20,22^. Functionally, through the utilization of real-time imaging technology, transplanted HSCs were often detected homing to the endosteal region, particularly to the N-cad^+^ niche within the TBA^24,25^. With the identification of improved HSC markers, Lin^-^ CD150^+^CD48^-^CD41^-^, the more accurate mapping the location of HSCs became possible as 1 out of 3 CD150^+^CD48^-^CD41^-^ cells were shown to support LT multilineage reconstitution in the repopulation assay^26^. Utilizing Immunofluorescent staining with Lin^-^CD150^+^CD48^-^CD41^-^ markers, HSCs were confirmed to localize in the endosteal niche (∼14-16%), but a large portion of these phenotypical HSCs (80-85%) were found within 5-cell distance from blood vessel, especially the sinusoid, thus the vascular niche concept was formed^26,27^. Subsequently, numerous groups have contributed to identification of various cell types that provide a niche function and these include Cxcl12-Aboundandt Reticular (CAR) cells, Nonmyelin Schwann cells, Nestin^+^, Prx1^+^, or Leptin receptor (LepR)^+^ stromal cells, as well as megakaryocytes (Mks)^7,27–33^. Given that the majority of stromal cells as described above are associated with vessels, and the main components of which are the sinusoidal structure primarily found within the CM covered by compact bone in the diaphysis region, the concept of the perivascular niche predominantly confined within the CM has emerged^5^. Consistent with the observation that 80-85% HSCs reside in the perivascular niche, deletion of *Cxcl12* or *KitL* (or SCF) from vessel and vessel-associated stromal, such as LepR^+^ or Prx1^+^, cells, led to a substantial (∼85%) loss of HSCs^6,7,34^. However, the location and characteristics of the remaining HSCs after the depletion of HSCs from the CM remained largely unclear. The following observation seems to provide an answer: myeloablation induced by the chemotherapy agent 5-fluorouracil (5-FU) resulted in a significant loss of HSCs primarily residing in perivascular niches, whereas the surviving, thus reserve, HSCs predominantly (∼75%) found in the endosteal niche within TBA^35^.

Among the cell types reported with niche functions, accumulating evidence strongly supports mesenchymal stem cells (MSCs) as pivotal in maintaining HSCs^28^. Several MSC markers have been identified within BM, including Nestin, Prx1, and LepR ^7,28,30^. While N-cad was the first identified niche marker in BM, the characteristic of N-cad^+^ stromal cells was not fully understood until they were revealed to possess the potential to differentiate into not only osteoblasts but also adipocytes and chondrocytes, thus validating N-cad as an MSC marker^35^. All the aforementioned studies shed light on niche markers and factors, providing valuable insights into understanding FL-HSCs and their interactions with niches.

While the FL primarily acts as a temporary site for HSC further maturation and expansion, numerous studies have focused on identifying the cellular and molecular components involved in supporting FL-HSC expansion. Previous investigations, primarily utilizing histological staining, *in vitro* culture, and single-cell RNA sequencing (scRNA-seq), have revealed complex interactions between FL-HSCs and various FL cell types, including ECs, hepatoblasts, hepatic stellate (stromal) cells, perivascular cells, and macrophages, mediated by diverse cytokines and chemokines^36–42^. Despite the undeniable roles of different FL structure cells and hematopoietic cells in supporting FL-HSCs, it still requires a systematic and comprehensive understanding of the niches responsible for FL-HSC maintenance and expansion.

Histological imaging, while capable of showing cell positions, often revealing only a limited set of markers. Single cell (sc)RNA-seq can provide an in-depth profile of gene expression but lacks the ability to preserve the spatial information of each cell within its original tissue context, making it challenging to comprehensively profile the HSC-niche interactions and signaling. In contrast, high-resolution spatial transcriptomics represents a promising and cutting-edge technology that integrates spatial information with transcriptomic data^43^. This approach enables the profiling of gene expression within their native spatial context, providing a powerful way to visualize gene expression patterns within a tissue or organ with unprecedented detail^43,44^. Hence, in this study, we employed high-resolution spatial transcriptomics, specifically the Slide-seq V2 method^45^, to generate a spatial transcriptomic cell atlas of the mouse FL. Additionally, we utilized genetic approaches to validate our observations derived from spatial transcriptomics. Our primary objective was to investigate the niches responsible for maintaining and expanding FL-HSCs.

## Results

**Single-cell RNA-seq reveals the transcriptomics of mouse fetal liver HSCs and potential niche cells.**

To establish a comprehensive cell atlas of HSCs and other cell types in FL, we conducted scRNA-seq of the mouse FL at E16.5. Given the rarity in the FL, we first isolated HSCs using fluorescence-activated cell sorting (FACS) and then combined them with CD45^-^ non-hematopoietic cells and CD45^+^ hematopoietic cells from the E16.5 FLs for scRNA-seq analysis, resulting in a total of 62,236 collected cells^46^. Subsequently, we employed the Uniform Manifold Approximation and Projection (UMAP) analysis to identify and classify 8 clusters of FL parenchymal cells and 22 clusters of hematopoietic cells (Fig. 1a). Through a differential expressed gene (DEG) analysis, we uncovered marker genes specific to HSCs as well as potential niche cells (Fig. 1b). Notably, we observed high expression levels of *Ifitm1*, *Hoxa7*, *Hoxa9*, *Cd27*, *Kit*, and *Ltb* in E16.5 FL-HSCs. Additionally, we found that well-known HSC surface markers, such as *Procr* (Epcr), *Ly6a* (Sca1), and *Slamf1* (CD150), were also enriched in E16.5 FL-HSCs (Fig. 1c). We examined gene expressions in various potential niche cell types, including MSCs, ECs, hepatoblasts/hepatocytes, megakaryocytes, and macrophages. FL-MSCs displayed enrichment of *Pdgfrα*, *Pdgfrβ*, *Acta2*, and *Fgfr1*. We then compared several known MSC markers in adult BM. *Nes (Nestin*) and *Cspg4* (encoding Ng2) were previously reported to be expressed in arterial pericytes that located in proximity to FL-HSCs and exhibited the ability to drive HSC expansion *in vitro*^38^. Our data showed that only a subset of FL-MSCs expressed with either *Nes* (39.87%) or *Cspg4* (Ng2) (10.66%). In fact, *Nes* was predominantly expressed in endothelial cells. Similarly*, Lepr* was predominantly enriched in endothelial cells^30^ and barely detected in MSCs (0.71%) (Fig. 1d). We discovered that *Cdh2* (N-cad) was highly expressed in MSCs (59.77%), an intermediate level in hepatoblasts, and low level in a subset of hepatocytes, thus displaying an expression pattern largely overlapping with that of *Cxcl12* in MSCs (75.00%) and hepatoblasts, but apart from hepatocytes (Fig. 1d,e). Additionally, other Niche factors, such as *Kitl* and *Igf2,* were found in both MSCs and hepatoblasts; but *Ptn and Thpo* were respectively detected in MSCs and hepatoblasts.

**Fig. 1.**
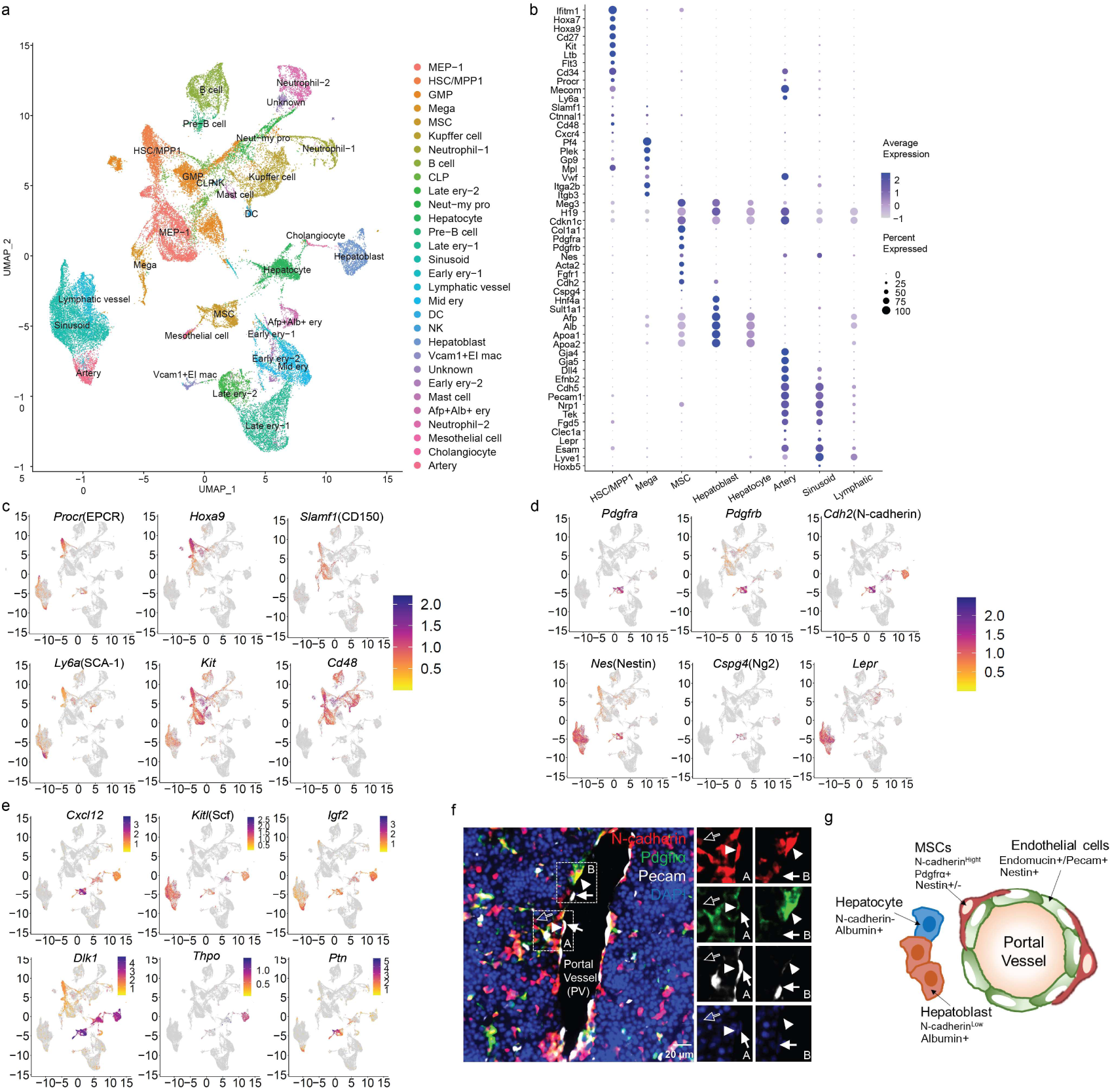
scRNA-seq profiling of E16.5 mouse FL reveals markers for FL-HSCs and potential niche cells. (a) UMAP visualization of E16.5 FL depicting distinct populations of HSCs and FL structure cells. (b) Dot plot representation displaying marker genes for HSCs and potential niche cells. (c) UMAP plot highlighting signature genes specific to HSCs. (d) UMAP plot showcasing signature genes specific to FL structure cells. (e) UMAP plot indicating the expression of niche factors in potential niche cells. (f) Immunofluorescence staining demonstrating the cellular architecture surrounding a portal vessel. Solid arrow: Pecam1^+^ endothelial cells; Solid arrowhead: N-cad^High^ Pdgfrα^+^ MSCs; Hollow arrow: N-cad^Low^ Hepatoblasts. Two peri-PV regions (A)(B) were highlighted and zoomed on the right panels. (g) Illustrations depicting the cellular architecture adjacent to a portal vessel.

We also conducted immunofluorescence staining to gain further insights into the cellular structure of the PV region of E14.5 FL. We identified four major cell types in the peri-PV region: inner layer comprised of Endomucin^+^/CD31^+^ ECs; the outer layer consisted of N-cad^High^Pdgfrα^+^ MSCs; and surrounded by N-cad^Int^ Albumin^+^ hepatoblasts. Notably, the majority of MSCs were observed adjacent to PV ECs in the peri-PV region (Fig. 1f, Extended Data Fig. 1a-d). For comparison, Nestin expression was detected in Endomucin^+^ ECs and a subset of N-cad^High^ MSCs (Extended Data Fig. 1e, f). Previously, FL stellate cells or perivascular mesenchymal cells with expression of Desmin and Acta2 were shown to play a role in supporting FL-HSCs^40^. In line with this report, our scRNA-seq and IF staining results revealed that N-cad^Hi^MSCs exhibited an enrichment of Desmin (Extended Data Fig. 1g, h). As a comparison, in the distal-to-PV (distal, hereafter) region, the hepatic sinusoidal was comprised of endothelial cells, hepatocytes, N-cad^Int^ hepatoblasts, and sporadically distributed N-cad^Hi^MSCs. The latter are not displayed as the perivascular pattern as seen in the PV region (Extended Data Fig. 1i).

In summary, we have effectively generated a fully annotated single-cell resolution atlas of the mouse FL, together with IF staining, we provided comprehensive insights into the cellular architecture of FL structure cells in the peri-PV region (Fig. 1g), and the sinusoidal structure at the distal region. Notably, our study unveils that N-cad-expressing cells constitute the primary source of niche factors in mouse FL.

### High-resolution spatial transcriptomics identifies HSCs and niche cells in mouse FL

To gain a comprehensive understanding of the interactions between HSCs and niche cells, we employed Slide-seq V2, a high-resolution (10µm) spatial transcriptomic technology in our study^45^. We collected two sections of E14.5 mouse FL, focusing specifically on a region enriched with portal vessels, as previous studies have suggested proximity of FL-HSCs to the PV structure (Fig. 2a, b, Extended Data Fig. 2a). Since Slide-seq had not been previously applied to mouse FL investigations, we conducted quality control analyses by assessing raw reads, high-quality reads, and exotic reads, and compared them with a previous study conducted on the mouse hippocampus^44^ (Extended Data Fig. 2b-e). Our Slide-seq data exhibited a median of approximately 320 unique molecular identifiers (UMI) per bead (d=10μm) and a median of around 240 genes per bead, with minimal reads corresponding to ribosomal genes (Extended Data Fig. 2i-l). Initially, using UMAP analysis, we identified only 10 distinct cell clusters due to the lower UMI and gene reads compared to scRNA-seq (Extended Data Fig. 2 f-g). To construct a comprehensive high-resolution spatial transcriptomic atlas of the E14.5 mouse FL, we applied the Robust Cell Type Decomposition (RCTD) algorithm to assign cell types to the beads based on expression profiles derived from E14.5 scRNA-seq^47^. This supervised approach facilitated the categorization of 28 cell types, including 24 hematopoietic cell types and 4 parenchymal cell types (Extended Data Fig. 2 h).

**Fig. 2.**
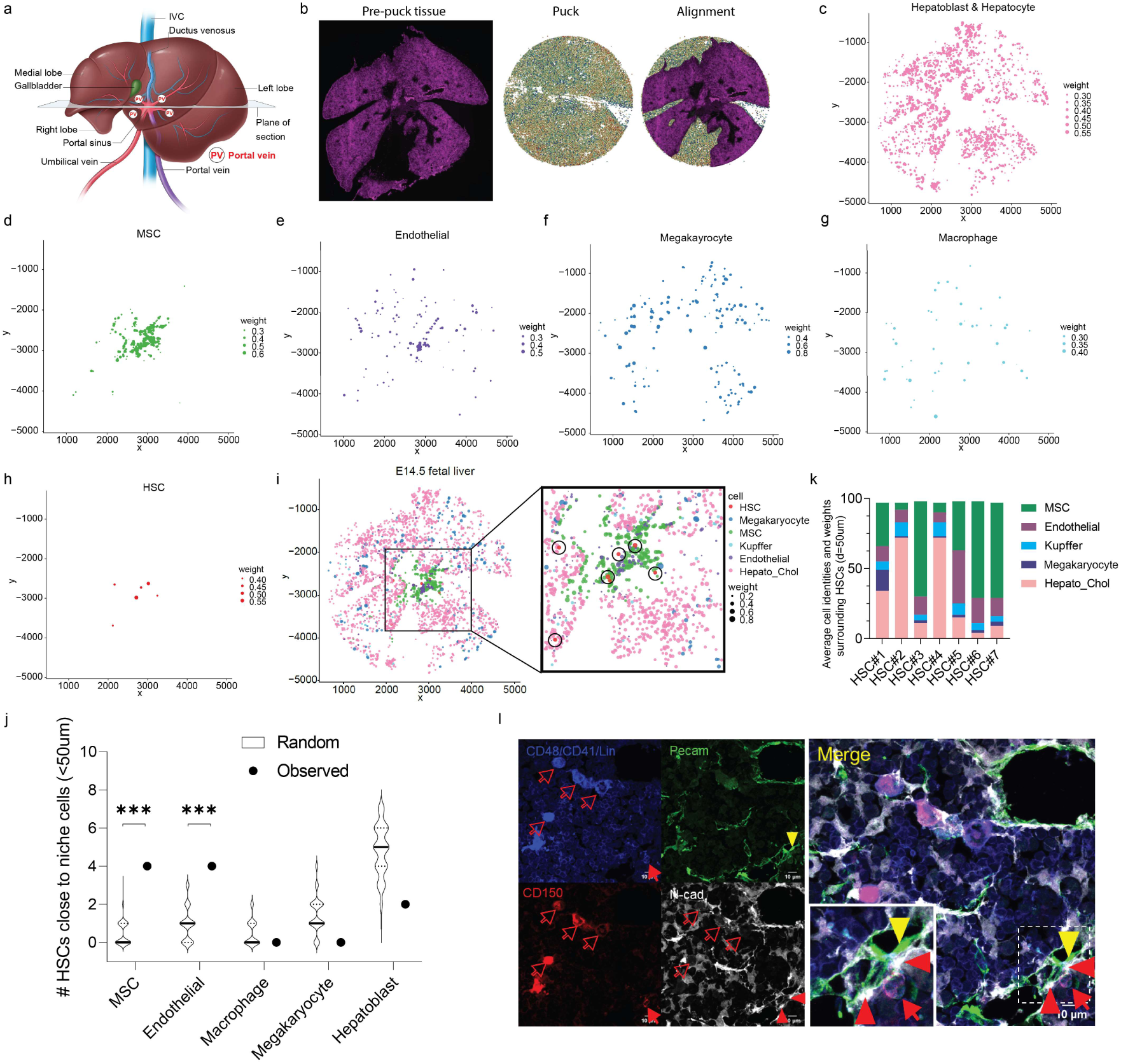
Mapping cellular architecture and interactions in E14.5 mouse FL using Slide-seq. (a) Illustrations depicting the mouse FL and plane of sections. (b) DAPI staining on pre-puck section (left), the reconstructed SLIDE-seq puck post-sequencing (Middle), and the alignment of the tissue on the puck (Right). (c)-(g) Identification of hepatoblasts & hepatocytes, MSCs, endothelial cells, megakaryocytes, and macrophages (Kupffer cells) using Slide-seq with a threshold of 25% (15% for endothelial cells). (h) Identification of HSCs with a threshold of 35%. (i) Cellular architecture of E14.5 mouse FL. Circles highlight the FL-HSCs. (j) Permutation test demonstrating the enrichment of HSCs to potential niche cells. The closest niche cell within 50μm to HSCs was considered as the niche cell to HSCs. (k) Bar graph presenting the average cell identities surrounding each HSC from puck#1 within 50μm. Decomposition was applied to all the beads surrounding each HSC within 50μm. The weights of erythrocytes and erythroblasts were excluded on all the beads. (l) Immunofluorescence staining illustrating the HSCs-MSC interactions. Solid red arrow: CD150^+^Lin^-^CD48^-^CD41^-^ HSC; hollow red arrow: CD150^+^LIN^+^CD48^+^CD41^+^ Hematopoietic progenitor and lineage cells; red arrowhead: N-cad^High^ MSCs; yellow arrowhead: Pecam1^+^ endothelial cells. ***, p<0.001. showing combined p-value calculated using Fisher’s combined probability test

We initially detected the presence of signature genes associated with erythrocytes and erythroblasts across the entire “puck”(a rubber-coated glass coverslip containing a monolayer of randomly deposited, DNA barcoded beads)^43^ (Extended Data Fig. 3a). As anticipated, the signature genes specific to the main structural and functional cells in the FL, such as hepatoblasts and their progeny hepatocytes, were predominantly detected within the tissue-covered regions (Extended Data Fig. 3b). To investigate the spatial distribution of FL cells and the interaction between HSCs and niche cells, we assigned the beads as credible representatives of a specific cell type when the enriched signature genes exceeded a certain threshold. For potential niche cells, we first annotated hepatoblasts and hepatocytes using a threshold of 25%, which best represented the spatial localization of these cell types on the puck (Extended Data Fig. 3c, d). Utilizing this threshold, we identified five potential niche cell populations on the pucks, each displaying its distinct spatial distribution pattern. Specifically, hepatoblasts and hepatocytes were major parenchymal cells in the liver responsible for supporting structural and metabolic functions (Fig. 2c). Furthermore, we observed an enrichment of MSCs in the PV region, consistent with our prior IF staining findings (Fig. 2d). Most megakaryocytes and macrophages were distributed within the liver lobes. ECs either resided on the border of the PV as PV-ECs or were dispersed within the lobe as sinusoid ECs, most of them are localized at the distal region (Fig. 2 e-g). A total of 7 HSCs were identified on puck #1, and 5 HSCs were identified on puck #2 using a threshold of 35% (Fig. 2h, Extended Data Fig. 3e). Previous studies have estimated that the total number of FL-HSCs is ∼500 at E14 and ∼1100 at E15, and FL-HSCs are enriched surrounding portal vessel^13,38^. Based on the thickness of our section (10um) and an average of 6 FL-HSCs identified per section, we estimated that the total number of FL-HSCs per E14.5 liver in our experiment is less than 1200, which is close to previous findings. It is noteworthy that the majority of HSCs were found in close proximity to MSCs within the PV regions (Fig. 2i, Extended Data Fig. 3f). Out of the 12 HSCs from both pucks, 7 were adjacent to MSCs, 5 were near ECs, 4 were close to hepatoblasts, 1 was in proximity to megakaryocytes, and none were near macrophages. To assess the significant enrichment of HSCs to niche cells considering the variable abundance of niche cell types in the FL, we performed a permutation test (see method). The results showed a significant enrichment of FL-HSCs near MSCs and ECs compared to other cell types (Fig. 2j, Extended Data Fig. 3g). Furthermore, to comprehensively explore the niche cells surrounding each HSC, we applied a decomposition method to identify the niche cell identities on each bead within a radius of 50μm or 30μm from each HSC. Our analysis consistently revealed that 8 out of 12 HSCs were located within a compartment enriched with MSCs, while 4 out of 12 HSCs resided in a compartment enriched with hepatoblasts and hepatocytes. In fact, 6 out of 12 HSCs tended to be in direct contact with MSCs. (Fig. 2k, Extended Data Fig. 4a-e). Immunofluorescence staining subsequently provided visual evidence of close spatial proximity between a Lin^-^CD48^-^CD150^+^ HSC and the N-cad^Hi^ MSCs as the outer layer of PV, as well as the CD31^+^ EC as the inner layer of PV (Fig. 2l).

To compare with a previous study that utilized the 10x Visium spatial transcriptomics solution for investigating FL-HSCs^41^, we employed the same method to three consecutive slices from an E14.5 mouse FL, following the standard protocol^41^. The 10x Visium method yielded an average of 5,139 median UMI per spot (d=55μm) and 1,756 median genes per spot, which were substantially lower than those obtained with SLIDE-seq (5,139 vs 320, d=55μm vs d=10μm; 1,756 vs 240, d=55μm vs d=10μm). Initially, the 10x Visium analysis identified nine cell clusters (Extended Data Fig. 5a, b). Consistently, MSCs were enriched in the PV region, exhibiting a gene enrichment score of approximately 80%, while megakaryocytes were predominantly located in the liver lobe region, with a gene enrichment score of around 30% (Extended Data Fig. 5c, d). However, the 10x Visium approach identified an average of two binding spots per section containing HSCs, with a gene enrichment score below 25% (Extended Data Fig. 5e). Similarly, to Slide-seq findings, the majority (4 out of 6) of HSC-containing binding spots were found adjacent to or overlapping with MSC-containing binding spots.

In summary, our high-resolution Slide-seq approach facilitated the generation of a comprehensive spatial transcriptomic map of the mouse FL at E14.5. In addition, the anatomical delineation of the PV structure through IF staining depicted MSCs and ECs respectively forming the outer and inner layers of PV. Collectively, our analyses unveiled a significant enrichment (>50%) of FL-HSCs within the PV region, positioned directly adjacent to MSCs and indirectly influenced by signaling emanating from ECs. While the enrichment of FL-HSCs to hepatoblasts didn’t reach statistical significance, the abundance of the latter suggests a potential interaction between them.

### Slide-seq decodes HSC-niche interactions

To investigate the dynamics of FL-HSCs and niche cells across various developmental stages, we performed unsupervised scRNA-seq clustering using a dataset comprising 50,804 cells obtained from FACS-sorted CD45^+^ and CD45^-^ populations at E14.5, E16.5, and E18.5 (Fig. 3a, Extended Data Fig. 6a). The differential expression of genes (DEG) analysis was conducted to identify representative genes of HSCs at the three embryonic time points. Notably, at E18.5, FL-HSCs exhibited a higher proportion of cycling cells, indicated by increased expression of *Mki67*, and decreased expression of *Cdkn1c (p57)*, *H19,* and *Mpl*. Previous studies have shown roles of *Cdkn1c (p57)*, *H19,* and *Mpl* in the maintenance of quiescent HSCs in the adult BM^48–50^. Additionally, we observed upregulation of FL-HSC signature genes such as *Procr*, *Hoxa9*, *Kit*, and *Mecom* during late gestation. However, genes associated with differentiation, including *Flt3*, and *Tgfβr1*, also showed increased expression levels. Consistently, we found a notable increase in the frequency of Pre-B and B cells, accompanied by a decrease in the frequency of HSCs and megakaryocytes from E14.5 to E18.5, which are consistent with the gradual increase of *Flt3* but decrease of *Mpl* and *Tek*^20,50,51^ (Fig. 3 b, c).

**Fig. 3.**
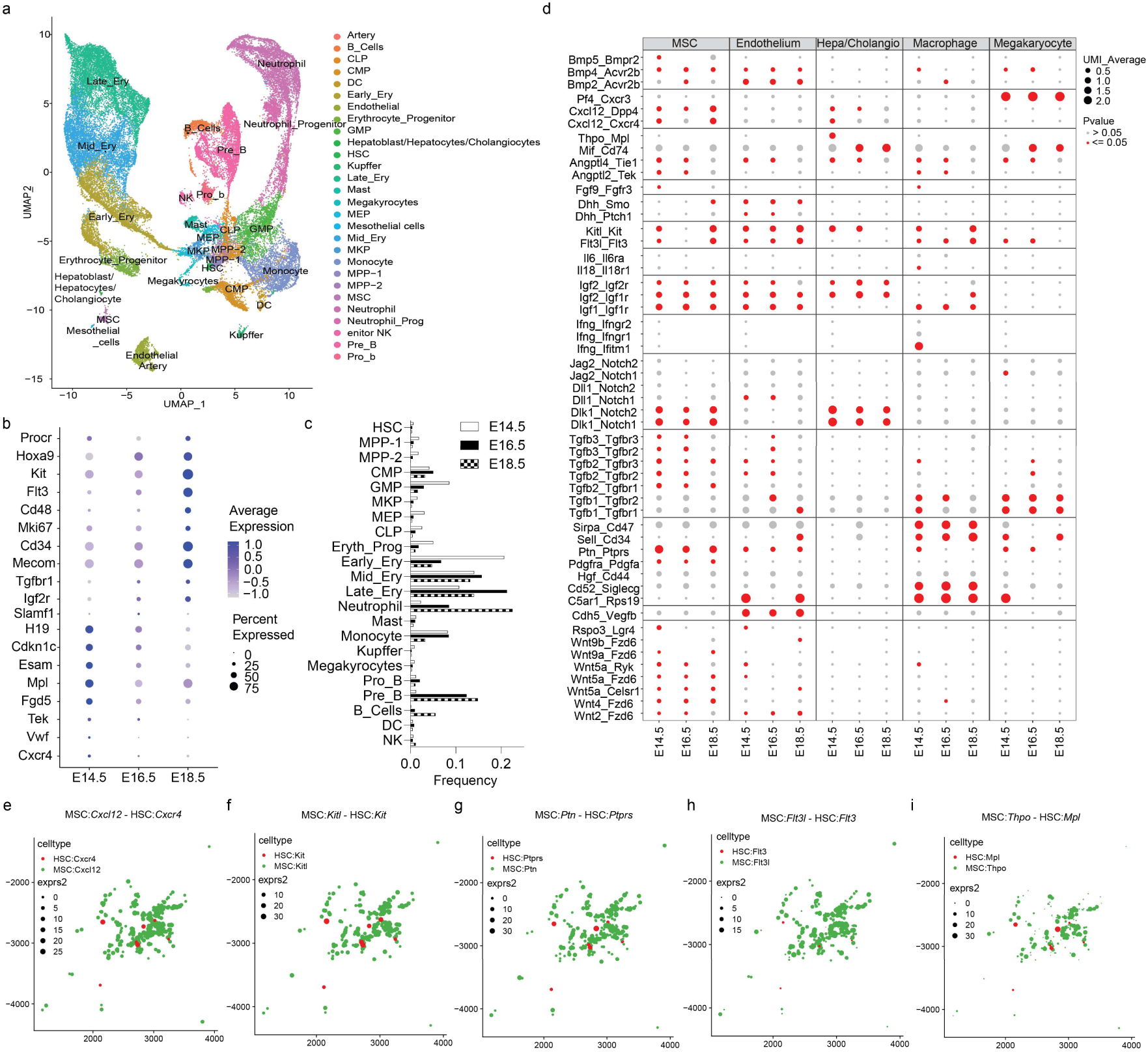
Predicting ligand-receptor interactions between niche cells and HSCs using Cellphone DB and SLIDE-seq. (a) Combined UMAP of mouse E14.5, E16.5, and E18.5 FL (b) Dot plot illustrating the changes in HSC signature genes across mouse E14.5, E16.5, and E18.5 FL. (c) Bar plot displaying the frequency of HSCs and lineage cells in mouse E14.5, E16.5, and E18.5 FL. (d) Cellphone DB analysis predicting niche-HSC signaling in E14.5, E16.5, and E18.5 mouse FL. (e) - (i) Slide-seq revealing the signaling interactions between MSCs and HSCs in E14.5 mouse FL.

To explore the potential ligand-receptor interactions between niche cells and HSCs in an unbiased manner, we utilized CellPhone DataBase (CPDB) analysis^52^. Specifically, we focused on characterizing interactions between HSCs and potential niche cells, including MSCs, ECs, hepatoblasts, megakaryocytes, and macrophages. Through comprehensive analysis, we identified MSCs as the primary providers of ligands involved in niche-HSC interactions. These included well-established niche signals known to support HSC in the adult BM, such as *Cxcl12-Cxcr4*, *Cxcl12-Dpp4*, *Igf1-Igf1r*, *Igf2-Igf2r*, *Dlk1-Notch1/2*, *Tgfβ*s*-Tgfβr*s, *Ptn-Ptprs*, *Kitl-kit*, and *Wnt* signaling pathways. Hepatoblasts shared most of the identified interactions with MSCs, apart from their unique enrichment in the *Thpo-Mpl* signaling module with HSCs, suggesting their role in supporting megakaryocyte fate. ECs shared several ligands with MSCs, including *Bmp4-Acvr2b*, *Kitl-Kit*, *Igf1-Igf2r*, *Igf2-Igf2r*, *Tgfβ2-Tgfβr2/3*, and *Ptn-Ptprs, but* lacked *Cxcl12-Cxcr4*, *Cxcl12-Dpp4*, and *Dlk1-Notch1/2* signaling modules. In fact, only N-cad-expressing cells, including MSCs and hepatoblasts, exhibited a *Cxcl12-Cxcr4* interaction, suggesting their role as major chemokine-releasing cells involved in HSC homing and localization. Macrophages and megakaryocytes displayed several distinct interactions respectively, including *Ifng-Ifitm1* and *Pf4-Cxcr3 respectively*, suggesting their unique contributions to support FL-HSCs (Fig. 3d).

While CPDB offers a scRNA-seq-based method for predicting ligand-receptor interactions between cell clusters, it lacks the ability to provide a spatial representation of genuine cell-cell interactions within an anatomically organized tissue. To verify the cell-cell interactions identified by CPDB analysis, we first employed a "projection" model to enhance genes detected on slide-seq (Extended Data Fig. 6b). Next, we assessed the "observed" expression profile of *Alb*, which ranked as the 3rd most enriched gene on the puck and compared it with the "projected" expression profile. The projection model effectively enhanced the expression level of *Alb* (Albumin) with high fidelity (Extended Data Fig. 6c-f). Similarly, the expression profile of *Cxcl12*, previously known to be enriched in MSCs, was significantly enhanced using the projection model (Extended Data Fig. 6 g, h). We then utilized Slide-seq to examine the spatial expression patterns of the ligand-receptor pairs predicted by CPDB analysis. Based on the number of HSCs involved in ligand-receptor interactions with niche cells, we defined these interactions as strong (involving at least 3 out of 7 HSCs, Extended Data Fig. 7a, b, f, j), weak (involving 1 or 2 out of 7 HSCs, Extended Data Fig. 7c, d, h, i), or no interaction (Extended Data Fig. 7e, j). Overall, MSCs exhibited several strong interactions with HSCs, including *Cxcl12-Cxcr4*, *Ptn-Ptprs*, *Kitl-Kit*, *Tgfβ2-Tgfβr2*, and *Dlk1-Notch1/2*. Additionally, MSCs displayed weak interactions such as *Cxcl12-Dpp4*, *Flt3l-Flt3*, *Igf1/2-Igfr1/2*, *Tgfβ1-Tgfβr1*, *Tgfβ3-Tgfβr3*, *Bmp4-Acvr2b*, *Bmp5-Bmpr2*, *Angptl2-Tie2*, *Wnt2-Fzd6*, and *Wnt5a-Celsr1/Fzd6/Ryk* (Fig. 3e-i, Extended Data Fig. 7k). These findings indicate that MSCs are a major source of niche factors that maintain FL-HSCs in proximity.

### The depletion of Cxcl12 from N-cad-expressing cells leads to increased frequency of HSCs exhibiting lineage-biased differentiation

It has been established that Cxcl12 serves a crucial niche factor for maintaining HSCs in the adult BM, demonstrated by studies showing that the loss of Cxcl12 from BM vessel and vessel-associated stromal cells resulted a large reduction of HSC frequency^7^. Given our observation that *Cxcl12-Cxcr4* interactions predominantly occur between HSCs and N-cad-expressing cells, including MSCs and hepatoblasts, we generated a conditional *Cxcl12* knockout mouse model using the *Cdh2^CreERT^;Cxcl12*^f/f^ strain to investigate the role of Cxcl12 in the maintenance of HSCs in the FL (Fig. 4a). Subsequently, we performed fluorescence-activated cell sorting assays to assess the impact of Cxcl12 loss on the frequency of HSCs, B cells, and CD41^+^ megakaryocytes (Extended Data Fig. 8a). Unexpectedly, conditional knockout of Cxcl12 from N-cad-expressing cells resulted in a significant (approximately 2-fold) increase in HSCs when normalized to CD45^+^ cells (∼1.5-fold increase normalized to total liver cells) (Fig. 4b). FL B cells were completely depleted upon loss of Cxcl12 (Fig. 4c). Conversely, the number of megakaryocytes increased significantly (∼2-fold) when normalized to CD45^+^ cells (Fig. 4d).

**Fig. 4.**
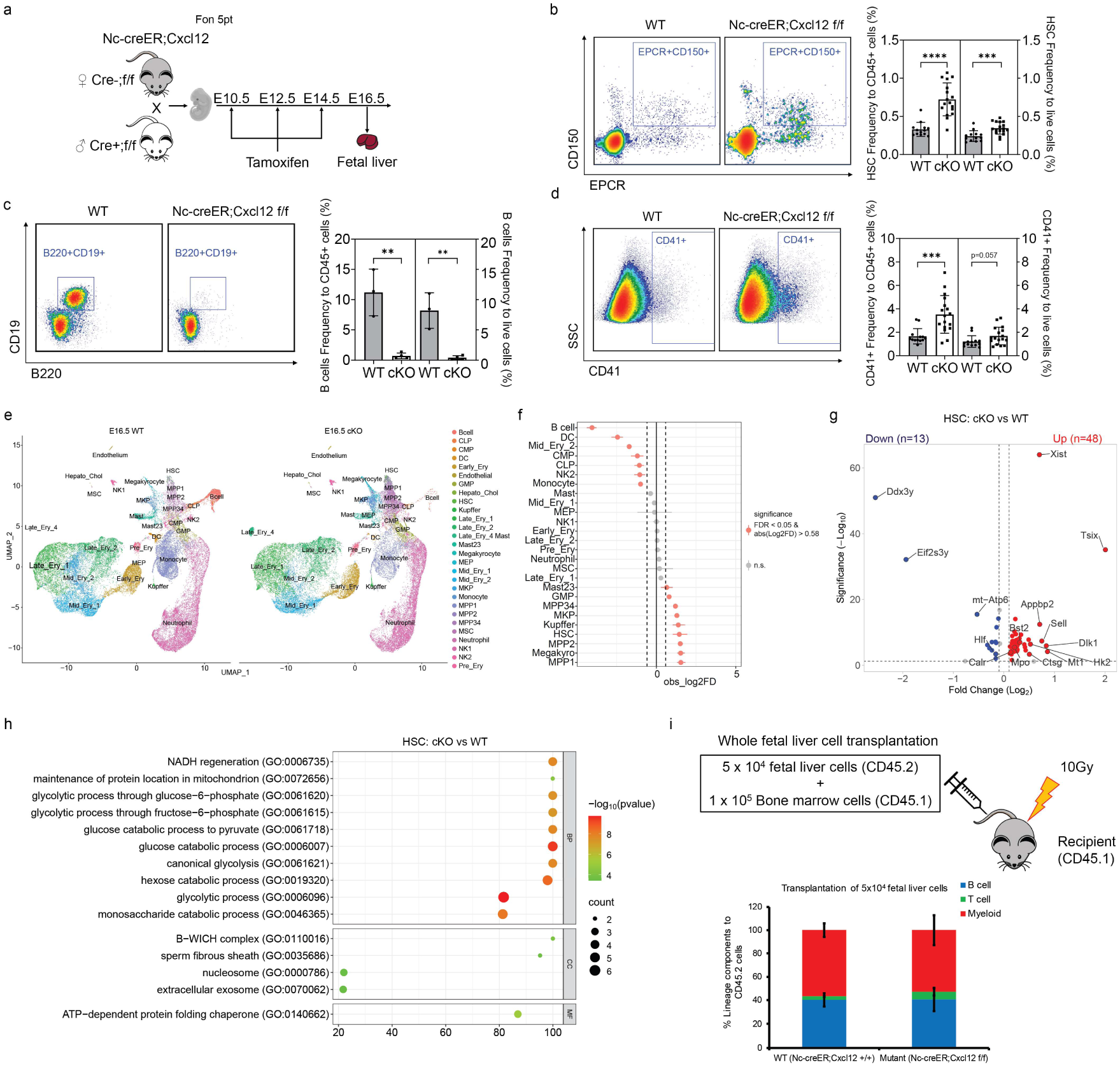
Conditional knockout of Cxcl12 from N-cad expressing cells leads to HSC expansion with myeloid bias. (a) Schematic overview of the experimental workflow. (b) - (d) Representative FACS profiles and bar plots demonstrating the expansion of HSCs (b), depletion of B cells (c), and increased megakaryocytes (d) following the loss of Cxcl12 from N-cad-expressing cells. (e) UMAP visualization of E16.5 mouse FL from wild-type (WT) and conditional knockout (cKO) mice. (f) Bar plot indicating the skewed myeloid differentiation after the loss of Cxcl12 from N-cad expressing cells. (g) Volcano plot displaying differentially expressed genes (DEGs) in HSCs mediated by Cxcl12 cKO from N-cad-expressing cells. (h) Bubble plot illustrating the top Gene Ontology (GO) terms in HSCs (BP: biological process; CC: cellular components; MF: Molecular Functions) affected by Cxcl12 cKO from N-cad-expressing cells. (i) Bar plot showing the frequency of B cells, T cells, and myeloid cells in the recipient BM 16 weeks post-transplantation. *, p<0.05; **, p<0.01; ***, p<0.001. showing p-values calculated using student t’s test.

Consistently, scRNA-seq analysis revealed that the loss of Cxcl12 had a significant impact on HSC proliferation as evidenced by increased HSC number and skewed lineage differentiation. Specifically, there was an increased myeloid cell such as megakaryocytes and their progenitors, Inversely, there were loss of common lymphoid progenitors (CLPs) and B lymphocytes upon conditional knockout of Cxcl12 from N-cad-expressing cells (Fig. 4e, f). DEG analysis showed that Cathepsin G (*Ctsg)*, *Mpo*, Selectin L (*Sell)*, and *Tetherin* were increased in HSCs after Cxcl12 loss, providing the molecular basis for an increase in HSC proliferation, mobilization, as well as an increase in differentiation towards myeloid cells ^53–56^ (Fig. 4g). Moreover, Gene ontology (GO) analysis revealed an enrichment of pathways related to NADH regeneration and glycolysis in HSCs upon Cxcl12 loss (Fig. 4h). N-cad-expressing cells, particularly MSCs together with ECs, hepatoblasts in the PV niche, are responsible for maintaining FL-HSCs with their full lineage potential. DEG analysis of MSCs revealed that MSCs in wild-type (WT) mice are enriched in *Igf1* and *Cxcl12*, indicative of their supportive role in HSC maintenance^57^ (Extended Data Fig. 8b). Subsequently GO analysis indicated that MSCs in WT mice are enriched in pathways related to extracellular matrix organization and collagen fibril organization, suggesting that extracellular remodeling by MSCs may create a favorable microenvironment with appropriate geometric and mechanical properties to harbor HSCs^58^ (Extended Data Fig. 8c). In contrast, MSCs in conditional knockout (cKO) mice lost the enrichment of extracellular matrix organization pathways, indicating an unfavorable extracellular matrix microenvironment for HSCs harbor (Extended Data Fig. 8d).

Based on the CPDB analysis, the predicted significant ligand-receptor signaling from MSCs to HSCs, including *Dlk1-Notch1/2*, *Ptn-Ptprs*, *Cxcl12-Cxcr4*, *Cxcl12-Dpp4*, and *Igf1-Igf1r*, were reduced following the cKO of *Cxcl12*. Of particular note, the *Flt3l-Flt3* signaling module between MSCs and HSCs was lost, most likely responsible for loss of CLP and downstream B cells^59^. Several other ligand-receptor signaling interactions between potential niche cells and HSCs were also found to be reduced, such as *Pf4-Cxcr3* and *C5ar1-Rps19* between Megakaryocyte and HSCs, *Igf1-Igf1r* and *C5ar1-Rps19* between Kupffer cells and HSCs, and *Cxcl12-Cxcr4* between hepatoblasts and HSCs. Furthermore, *Tgfβ* signaling was predicted to be increased in hepatoblasts/ hepatocytes and Kupffer cells, while *Thpo-Mpl* signaling was increased in hepatoblasts/hepatocytes, suggesting their involvement in skewed myeloid differentiation^60,61^. Moreover, endothelial cells showed an increase in *Sirpa-Cd47* signaling, implying the potential involvement of immune privilege pathways to safeguard FL-HSCs against macrophage clearance and potentially contribute to HSC expansion^62^ (Extended Data Fig. 8e).

To further investigate the functional significance of Cxcl12 originating from N-cad-expressing cells in HSC maintenance, we isolated FL-HSCs from both WT and *Cxcl12*-cKO mice and transplanted them into WT adult recipient mice, which possess a normal expression of Cxcl12 from N-cad-expressing cells. Following 16-week period, we observed that the recipient mice exhibited balanced lineage reconstitution, indicating that the observed lineage bias resulted from *Cdh2^CreER^* induced *Cxcl12* depletion in the FL was a niche-dependent phenotype (Fig. 4i).

### The absence of Cxcl12 from N-cad-expressing cells results in the altered localization of HSCs to distinct niches

As Cxcl12-Cxcr4 is a recognized chemoattractant factor, we anticipated that the loss of Cxcl12 from N-cad-expressing cells including mainly MSCs and hepatoblasts (Fig. 1d) would lead to an altered localization of of HSCs. To test this, we employed Slide-seq using two FL sections from E16.5 WT and *Cxcl12*-cKO mice, respectively. Compared to the HSC distribution pattern in E14.5 mouse FL, although distribution of HSCs increased in the distal region, ∼50% of HSCs are still in the PV region and located close to MSCs in E16.5 FL (Fig. 5a). Intriguingly, depletion of Cxcl12 from N-cad^+^ cells resulted in the relocation of HSCs initially located at the PV niche to the distal region of fetal live (Fig. 5b). Specifically, in WT mice, 50% ± 23% of 36 HSCs were found within 50μm of MSCs, whereas, in *Cxcl12*-cKO mice, only 13% ± 3% HSCs were in proximity to MSCs. Similarly, in WT mice, 73% ± 9% of HSCs were close to ECs, compared to 38% ± 9% HSCs in *Cxcl12-*cKO mice (Fig. 5c, d). To assess the significance of HSC enrichment in proximity to various niche cells, we conducted a permutation test. The data revealed that HSCs in WT mice were significantly enriched in proximity to MSCs and ECs, whereas HSCs in *Cxcl12-*cKO mice did not show such enrichment. Although the frequency of HSCs in proximity to ECs decreased after Cxcl12 knockout, it did not change significantly between WT and cKO mice (Fig. 5e, Extended Data Fig. 9a). These findings suggest that the relocation of HSCs away from MSCs and remain interaction with ECs at the distal regions, where enriching sinusoids (Fig. 1f). It is possible that hepatocytes remain relatively higher Cxcl12 level (Fig.1e) in mutant FL when Cxcl12 was depleted from N-cad relatively higher expressed MSCs and hepatoblasts (Fig.1d), which however requires a verification.

**Fig. 5.**
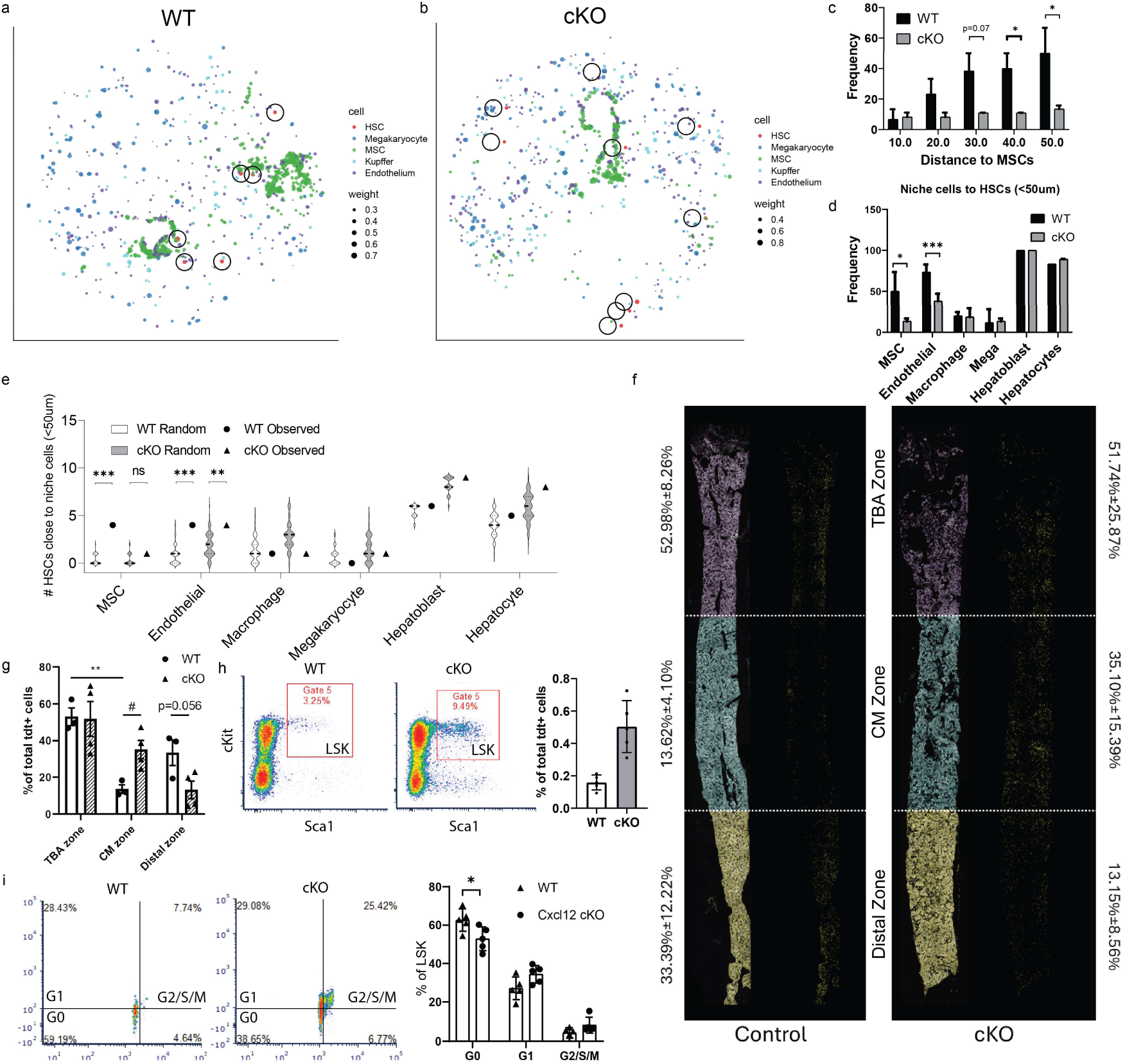
Altered localization of HSCs in both FL and adult BM after conditional knockout of Cxcl12 from N-cad-expressing cells. (a) (b) Slide-seq images displaying the cellular architecture in E16.5 mouse FL from wild-type (WT) (a) and conditional knockout (cKO) (b) mice (hepatoblasts and hepatocytes not shown). (c) Bar plot illustrating the distances between HSCs and MSCs in WT and cKO mice. (d) Bar plot indicating the significance of HSCs located within a distance of <50μm to niche cells in WT and cKO mice. (e) Permutation test demonstrating the enrichment of total HSCs in proximity to potential niche cells in WT and cKO mice. (f) Immunofluorescent staining of td-tomato showing the distribution of donor bone marrow cells (from mT/mG mice) in the recipient bone marrow after 8 weeks. Left images showing the segmentation of the bone (TBA zone vs. CM zone vs. Distal Zone). Right images showing the td-tomato+ donor cells ausing anti-td-tomato IF staining. (g) Bar plot indicating the distribution of td-tomato+ donor bone marrow cells in the three different regions of recipient bone marrow. (h) FASC analysis and bar plot showing the proportion of LSK cells in recipient bone marrow in control vs cKO mice. (i) FASC analysis and bar plot showing the cell cycle analysis of LSK cells in recipient bone marrow in control vs cKO mice. *, p<0.05; **, p<0.01; ***, p<0.001 (c)(d)(g)(h)(i) showing p-value calculated using student t’s test. (e) showing combined p-value calculated using Fisher’s combined probability test (g) showing p-value calculated using one-way ANOVA test. #,p<0.05, (g) showing p-value calculated using student t’s test.

Since FL-HSCs predominantly undergo proliferation, we further subclustered the E16.5 HSCs into cycling and non-cycling populations based on their gene expression profiles from scRNA-seq (Extended Data Fig. 9b-d). The non-cycling HSCs exhibited high levels of *Cdkn1c (p57)*, *Ctnnal1(α-catenin1)*, and *H19*, indicative of their quiescent state. We then mapped the location of cycling and non-cycling HSCs in the E16.5 FL (Extended Data Fig. 9e-h). The frequency of cycling HSCs was increased upon Cxcl12 loss, which is consistent with the expansion of HSCs (Extended Data Fig. 10a).

Our findings indicate that the PV niche consisting of MSCs, ECs, and potentially hepatoblasts in mouse FL plays a crucial role in maintaining relatively non-cycling HSCs with full lineage potential. The loss of Cxcl12 from N-cad-expressing cells leads to the altered location of FL-HSCs towards a distal sinusoidal niche, which comprises ECs and potentially abundant hepatocytes but reraly MSCs (Extended Fig. 10d). Occasionally, megakaryocytes and macrophages can also be part of sinusoidal niche. The sinusoidal niche is pivotal in supporting proliferation of HSCs, albeit with a myeloid-lineage bias.

### Dpp4^+^ but not Dpp4^-^ HSPCs predominantly homing to the trabecular bone area in adults

We have focused on the analyses of the interaction between MSCs and FL-HSCs through the Cxcl12-Cxcr4 axis, the CPDB analysis also revealed the interaction between MSCs and HSCs involving Cxcl12 and Dpp4 (CD26) in mouse fetal liver (Fig. 3d). Dpp4 is a cell-surface protease known to cleave and deactivate various peptide hormones and chemokines, including Cxcl12 as the top-ranked target, thereby reducing its biological activity. This interaction can significantly influence the chemotactic function of Cxcl12, impacting cell migration, mobilization, and homing of stem cells^63,64^. However, it is difficult to study HSC-membrane associated Dpp4 as there is neither known HSC-specific Cre mouse line nor robust HSC transplantation assay system at the fetal stage.

To investigate Dpp4’s role as a modulator in HSC migration and homing, we transplanted 5x10^5^ total nucleated BM cells from Dpp4 WT (Dpp4^+/+^-RFP) or Dpp4 KO mT/mG mice (Dpp4^-/-^-RFP) into WT recipients. Our findings indicated that Dpp4-WT donor cells are notably enriched in the TBA region. Conversely, Dpp4-KO cells showed a relatively lesser-extent and exhibit a more evenly distributed pattern (Extended Fig. 11a, b). Furthermore, Dpp4-WT cells localized closer to N-cadherin+ cells at the endosteal niche compared to Dpp4-KO cells (Extended Fig. 11c).

Given Dpp4’s inhibitory effect on the Cxcl12-Cxcr4 axis due to its enzymatic activity, we hypothesized the presence of another factor in the endosteal niche modulating Dpp4’s activity. Previous studies have showed Glypican-3 (Gpc3) as an inhibitor of Dpp4 function^65,66^. Indeed, we observed that enriched Gpc3 correlated with Cxlc12 protein level in the endosteal regions of TBA in mouse BM, presumably though inhibiting Dpp4 and enhancing the Cxcl12-Cxcr4 signaling (Extended Fig. 11d). In contrast, the central marrow has decreased level of Gpc3 but shows high levels of Dpp4 expression primarily in CD31^+^ endothelial cells (Extended Fig. 11e). The combination of lower Gpc3, along with elevated Dpp4, in central marrow contributes to the relatively lower gradient of Cxcl12 in CM compared to TBA, thus hindering HSC homing.

### The absence of Cxcl12 from N-cad-expressing cells results in the altered localization of BM HSPCs from the trabecular bone region to the central marrow region

To further investigate how the Cxcl12-Cxcr4 axis influences HSC homing to the N-cadherin+ niche, we conducted a bone marrow transplantation assay in adult mice. This approach was chosen because transplantation into fetal mice presents significant challenges. We transplanted 1x10^6^ total nucleated bone marrow cells isolated from mT/mG mice into WT recipient (n=6), or the recipient with conditional knockout (cKO) of Cxcl12 from N-cadherin-expressing cells (n=5). After 8 weeks, the femurs were collected from both groups and those with intact BM on paraffin embedded sections were used for IF staining of RFP to determine the distribution of transplanted donor cells. The distribution of transplanted donor cells was significantly enriched in the trabecular bone area (TBA, the knee joint side) in WT recipients, compared with the central marrow (CM) area (TBA vs. CM, 52.98% ±8.26% vs. 13.62±4.10%). Upon conditional knockout of Cxcl12 from N-cadherin-expressing cells, transplanted donor cells increased homing to the CM from 13.62% (Wt) to 35.10% (mutant) (Fig. 5f, g). FASC analysis of the bone marrow from both groups showed that the LSK cell number was significantly increased in cKO mice compared to WT mice (9.49% vs 3.25%), indicating that CM niche supports the proliferation of HSPCs (Fig. 5h). We further conducted cell cycle analysis on the LSK cells in both groups. Data showed that in WT mice, in which the majority of HSPCs reside in TBA area, 59.19% LSK cells are in G0 phase, 28.43% LSK cells are in G1 phase, and 7.74% LSK cells are in G2/S/M phase. As a comparison, LSK cells in cKO mice exhibited a more proliferative profile, with 38.65% LSK cells are in G0 phase, 29.08% LSK cells are in G1 phase, and 25.42% LSK cells are in G2/S/M phase (Fig. 5i).

## Discussion

The journey of identifying and characterizing niches supporting HSCs has been lengthy and intricate, marked by significant progress but often filled with conflicting findings over the past decades. This complexity stems from both the complicated nature of the BM system, and the influence of the single cell-based niche model, resulting in a complex and controversial understanding of cell types crucial for HSC maintenance and function^1–5^.

The first niche discovered as supportive of HSCs in the adult mouse BM was the endosteal niche, situated within the TBA^18,19^. This niche, composed of osteogenic lineage or N-cad-expressing bone lining cells that now are recognized as MSCs^67^, was found to support LT quiescent HSCs^18,20,67^. Functional assays demonstrated the crucial role of the endosteal niche in attracting homing of and supporting restoration of HSCs in response to severe stress^35^. This aligns with the notion that the endosteal niche predominantly supports a ‘dormant’ (deep quiescent) state of HSCs, functioning as a reserve pool^18,22,68^.

Subsequent studies uncovered additional HSC niches in the adult BM, including the vascular niche, occupying 25% of the BM volume^69^ and hosts 80-85% of HSCs^26,27^. Mature sinusoidal vessels, covered by compact bone in the diaphysis region, associate various cell types in the perivascular region, contributing to niche functions^7,20,27–31,67,70,71^. Deletion experiments targeting molecules like Cxcl12 or Kitl from these vessels and vessel-associated MSCs, or myeloablation using 5-FU, resulted in a significant (80-85%) reduction of HSCs, underscoring the importance of these niches in supporting HSCs during homeostasis^6,7,34,35^.

In contrast to the adult BM, FL-HSCs were thought to have unique proliferative features, suggesting a distinct niche microenvironment^72^. Prior studies revealed potential niche cell types in FL, including arteriolar ECs, PV-associated stroma cells, and macrophages^38,41,42^. In this study, high-resolution spatial transcriptomics uncovered existing two niches within different zones of FL: the PV niche, maintaining quiescent HSCs with full lineage potential, and the sinusoidal niche, supporting proliferative HSCs with a bias towards myeloid lineage. Loss of Cxcl12 from N-cad-expressing niche cells altered HSC locations, leading to an “aging” phenotype with increased proliferation but with myeloid bias.

These findings refine our understanding of FL-HSC niches and provide insights for adult BM. Indeed, depletion of Cxcl12 from N-cad^+^ endosteal niche in adult BM altered the location of HSPCs from initially being in the TBA to the CM, resulting in a reduction in the G0 fraction of HSPCs. This is akin to the initial observation that Cxcr4^-/-^ mice exhibited a decreased G0 fraction of HSPCs, with HSPCs predominantly found in the CM, leading to the previous conclusion that Cxcl12-Cxcr4 signaling induces quiescence of HSPCs ^27^. However, in this study, with Cxcl12 lost from N-cad-expressing cells mainly in TBA, Cxcl12-Cxcr4 signaling between relocated HSPCs and niche cells remained intact in CM, the increased HSPC proliferation is due to microenvironmental differences between TBA and CM, rather than Cxcl12-Cxcr4 signaling *per se*. This revised interpretation offers a plausible explanation for why only Prx1^Cre^ but not Lepr^Cre^ mediated Cxcl12 deletion resulted in substantial HSPC loss in the BM and a significant increase in extramedullary hematopoiesis, despite overlapping expression patterns of Prx1 and Lepr in the BM^6,7^. This discrepancy is attributed to Prx1 initiating expression in the fetal stage, while Lepr is expressed only in adult^73^. Thus, Prx1^Cre^ but not Lepr^Cre^ mediated Cxcl12 deletion altered the Cxcl12 gradient between the BM and extramedullary hematopoietic sites at the fetal-adult transition stage, affecting neonatal homing of HSPCs to BM and subsequently reducing adult BM HSPCs. Similarly, this also account for why Ng2^Cre^ mediated, but not Ng2^CreEr^ induced, Cxcl12 deletion led to a decrease in the G0 fraction of HSPCs^74^. Our study, along with previous findings, has led us to propose a new model featuring two distinct niches with multiple cellular components in different zones. The TBA includes N-cad^+^ MSCs, osteogenic cells, various ECs, neural cells, supporting deep-quiescent HSC^75,76^. The CM includes sinusoidal ECs, vessel-associated MSCs, Schwann cells, and MKs, supporting heterogenous including relatively active HSPCs^7,20,27–31,67,70,71^. TGFβ1 from MKs, activated by Schwann cells, maintains a short-term quiescent state of HSCs during homeostasis but supports HSC restoration post BM injury via FGF signaling ^29,31,33,61^. Imaging of myelopoiesis defined the anatomy of normal and stress hematopoiesis near megakaryocytes^77^.

In summary, spatial transcriptomics has significantly advanced our understanding of HSC niche interactions. The new discovery about the interplay between Dpp4 and Gpc3, which plays a critical role in establishing the gradient of Cxcl12 across FL and BM, enhances our understanding of how the broad expression of Cxcl12 leads to the specific homing of HSPCs in both the FL and BM. This comprehensive analysis combining spatial transcriptomics, immunofluorescent imaging, and genetic approaches, results in a paradigm shift regarding the niche concept, moving from single cell-based to multicellular components within distinct zones as proposed previously^78^. This shift reflects the intricate regulation of HSPCs in association with their cycling states and metabolic activities according to the hematopoietic stages and in response to varies stresses, offering valuable insights for hematopoietic homeostasis, aging process, and leukemogenesis.

## ACKNOWLEDGMENTS

We thank K. Ferro, K. Bae, F. Liu, Z. Yu, E. Hamilton, M. Treese, I. Dunwiddie, J. Morrison, and M. Peterson, M. Miller for technical support. We would like to acknowledge the Board Institute and Dr. Fei Chen’s group for providing the Slide-seq V2 pucks. We would like to acknowledge the generated data on the Illumina NovaSeq 6000 System at University of Kansas Medical Center’s Genomics Core, which was supported by following grants: NIH U54 HD 090216, P30 GM122731-03, and NIH S10OD021743. This work was supported by the Stowers Institute for Medical Research (SIMR-1004) and NCI-designated Kansas University Comprehensive Cancer Center (P30 CA168524).

## AUTHOR CONTRIBUTIONS

R.D. conceived the project, designed, and performed experiments, analyzed data, and wrote the manuscript. H.L and J.R. performed bioinformatics analysis. R.D., C.W., W.L., S.M., K.P., K.H., A.S, M.M., S.H., and Z.Y. performed scRNA-seq, Slide-seq, immunofluorescence staining, 10X Visium experiments, and imaging processing and analysis. X.C.H. and M.H. provided the training. W.L., X.M., Z.Y., and M.E. provided technological support. M.E. performed proof-reading of the manuscript. S.S., S.M., Y.W., A.P., J.H., J.U., and B.S. provided technical assistance. X.K., X.C.H. and L.L. supervised the study and revised the manuscript.

## Method (main text)

### Mice

The mice utilized in this study were all housed in the AAALAC-accredited animal facility located at the Stowers Institute for Medical Research (SIMR), and were handled according to both SIMR and National Institutes of Health guidelines. All procedures involving experimental animals were approved and authorized by the Institutional Animal Care and Use Committee (IACUC) of SIMR. N-cad-CreER^T^ mice were generated by Applied StemCell, Inc. To induce N-cad-CreER^T^; Cxcl12 f/f mouse, Tamoxifen (Sigma) was injected intraperitoneally to the pregnant mothers at 2mg per injection at E10.5, E12.5, and E14.5. C57BL/6, Ptprc, and Cxcl12 floxed mice were acquired from the Jackson Laboratory.

### Tissue preparation and handling

FL tissue was received freshly harvested in 1x PBS and were individually submerged in OCT to remove excess PBS. Tissues were moved to a cryo-mold with fresh OCT and orientated in “tulip” orientation for transverse tissue sectioning. Tissue blocks were then flash frozen at -70°C before sectioning (HistoChill, Novec™ 7000). Region of interest (multiple portal/central vein openings on same plain) were identified prior to tissue capture.

### ScRNA-Seq

Dissociated cells in PBS + 2.5% FBS were assessed for concentration and viability via a Luna-FL cell counter (Logos Biosystems). Cells deemed to be 75-98% viable were loaded on a Chromium Single Cell Controller (10x Genomics), based on live cell concentration. Libraries were prepared using either the Chromium Single Cell 3’ Reagent Kits v3 or the Chromium Next GEM Single Cell 3’ Reagent Kits v3.1 (10x Genomics) according to manufacturer’s directions. Libraries were sequenced to a depth necessary to achieve at least 18,000 mean reads per cell on an Illumina NovaSeq6000 or Illumina NextSeq500.

### Slide-seq V2

The Slide-seq pucks used in this study were generated and sequenced at the Broad Institute (Cambridge, MA) by Dr. Fei Chen’s group according to the methods and Extended Data information provided in the Nature Biotechnology publication “Highly sensitive spatial transcriptomics at near-cellular resolution with Slide-seq V2” ^44^. The spatial barcode sequencing file of the pucks were generated via monobase ligation chemistry and provided by Dr. Chen’s group to be used for later analysis. The tissue was sectioned to 10 µm thickness and mounted directly onto the pucks using a cryostat (CryoStar NX70). Subsequent library preparation was performed according to manufacturer’s directions for the Nextera XT kit (Illumina, FC-131-1096)

### 10x Visium

Four 10um-thick tissue sections were adhered to the barcoded spots within the four individual capture areas of a 10x Genomics Visium Spatial Gene Expression Slide. The tissue was permeabilized for 18 minutes then processed to create cDNA and next generation sequencing libraries according to manufacturer’s instructions for the Visium Spatial Gene Expression Reagent and Slide Kits (10x Genomics, PN-1000189 and PN-1000188). Libraries were pooled at equal molar concentrations and sequenced to a depth necessary to achieve at least 50,000 read pairs per tissue covered spot on the Illumina NextSeq 500 instrument using a High Output flow cell.

### Flow Cytometry

Cells from mouse FLs or recipient BM freshly collected in 1xPBS with 2.5% PBS. Cells were then stained with surface markers for 30 min at 4°C and washed with PBS (with 2.5% FBS) trice. Cells were then stained with 7-ADD/DAPI for 5 mins on ice and then analyzed by ZE5 (Bio-Rad) or sorted by S6 cell sorter (BD Biosciences). FAC express software 7.0 was used for data analysis.

For cell cycle analysis, BM cells were fixed in 4% PFA at room temperature for 30min, followed by surface marker staining for 30 min at 4°C. Cells were then incubated with anti-Ki67 in 1xPBS+0.1% tween-20 overnight at 4°C, followed by DAPI staining for 5 min on ice and then analyzed by ZE5 (Bio-Rad).

Fluorescence-activated cell sorting was performed on S6 cell sorter (BD Biosciences) for the enrichment of HSC and niche cells for scRNA-seq. Cd150^+^, EPCR^+^, N-cad^+^, CD31^+^ cells were collected and pooled with CD45^+^ and CD45^-^ cell for scRNA-seq.

### Bioinformatics and Statistical analysis

To determine whether certain niche cell types are in the same niche micro-environment and in proximity to HSCs, we applied a permutation test. First, a fixed number equal of beads are randomly selected from all beads for a given puck and annotated as “pseudo” HSCs. The number of the “pseudo” HSCs are equal to the number of “real” HSCs identified on each puck. Second, beads and their annotated cell types within a 50um radius centered to the “pseudo” HSCs are identified. Finally, the number of niche cells are counted with each of these micro-environments (50um circle). This process was repeated 100 times, and the proportion of the number of nearby niche cells that are more extreme than our observed one is calculated. This proportion is the p-value showing how often the HPSCs and niche cells are in proximity.

See Extended Data method for the full bioinformatics analytical methods.

