## Supplementary figures for "Characterization of Multicellular Niches Supporting Hematopoietic Stem Cells Within Distinct Zones"

### Extended Data Figures

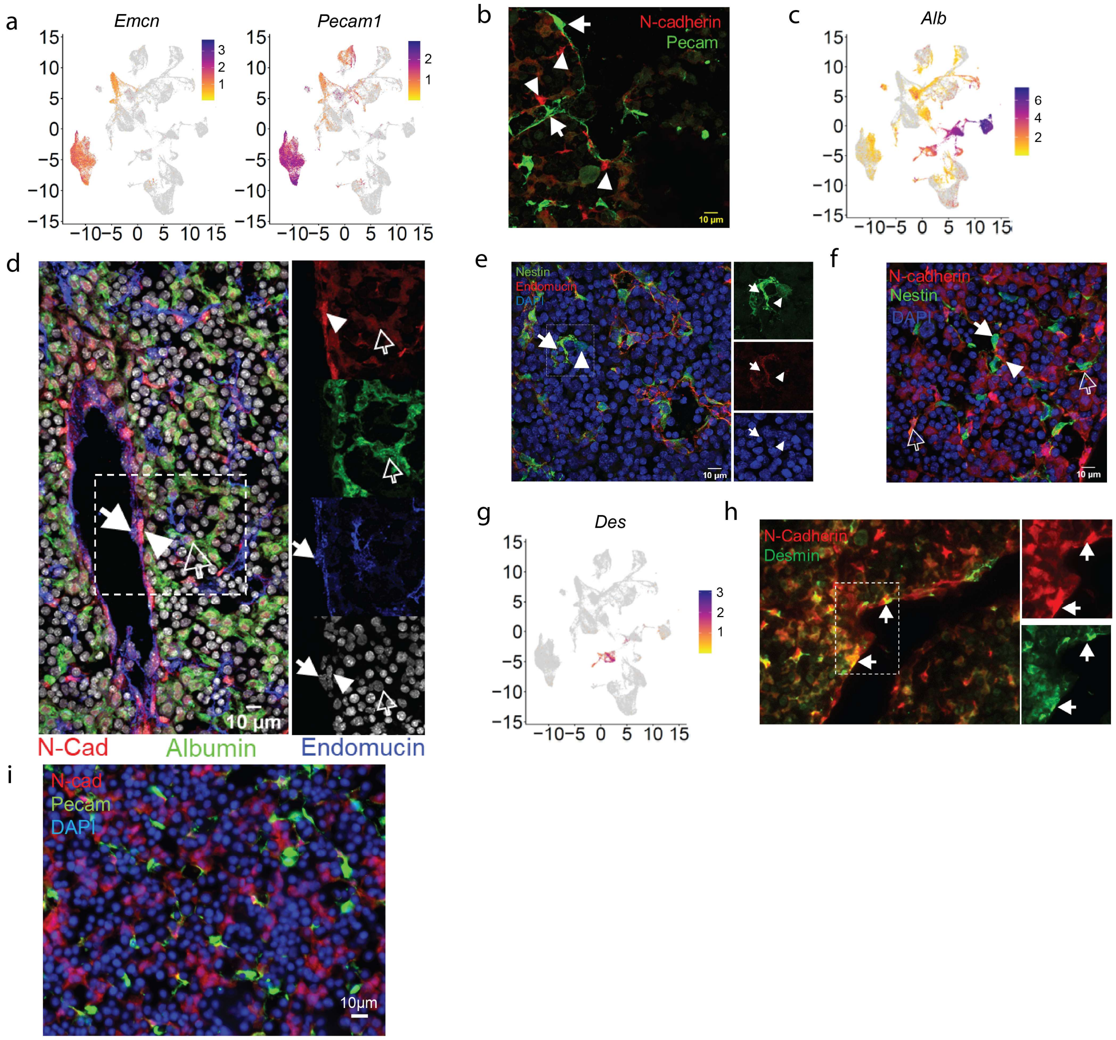


**Extended Data Figure 1. scRNA-seq and Immunofluorescent staining showing the cellular architecture surrounding portal vessels.**

(a) UMAP plot highlighting the expression of *Emcn* and *Pecam1* as endothelial cell markers.

(b) Immunofluorescence staining demonstrating N-Cad$^{High}$ MSCs are peri-portal vessel cells surrounding Pecam$^{+}$ endothelial cells. Arrow: Pecam$^{+}$ endothelial cells; Arrowhead: N-Cad$^{High}$ MSCs.

(c) UMAP plot highlighting the expression of *Alb* as hepatoblasts and hepatocyte markers.

(d) Immunofluorescence staining demonstrating the cellular architecture surrounding a portal vessel. Solid arrow: Endomucin$^+$ endothelial cells; Hollow arrow: Albumin$^+$ N-cad$^{Low}$ hepatoblasts; Arrowhead: N-cad$^{High}$ MSCs.

(e)(f) Immunofluorescence staining demonstrating the partial overlap of Nestin with Endomucin$^+$ endothelial cells (e), and N-cad$^{High}$ MSCs (g).

(g) UMAP plot highlighting the expression of Desmin as a liver stellate cell marker.

(h) Immunofluorescence staining demonstrating the overlap of Desmin with N-cad$^{High}$ MSCs.

(i) Immunofluorescence staining of N-cadherin and Pecam in the sinusoidal region.

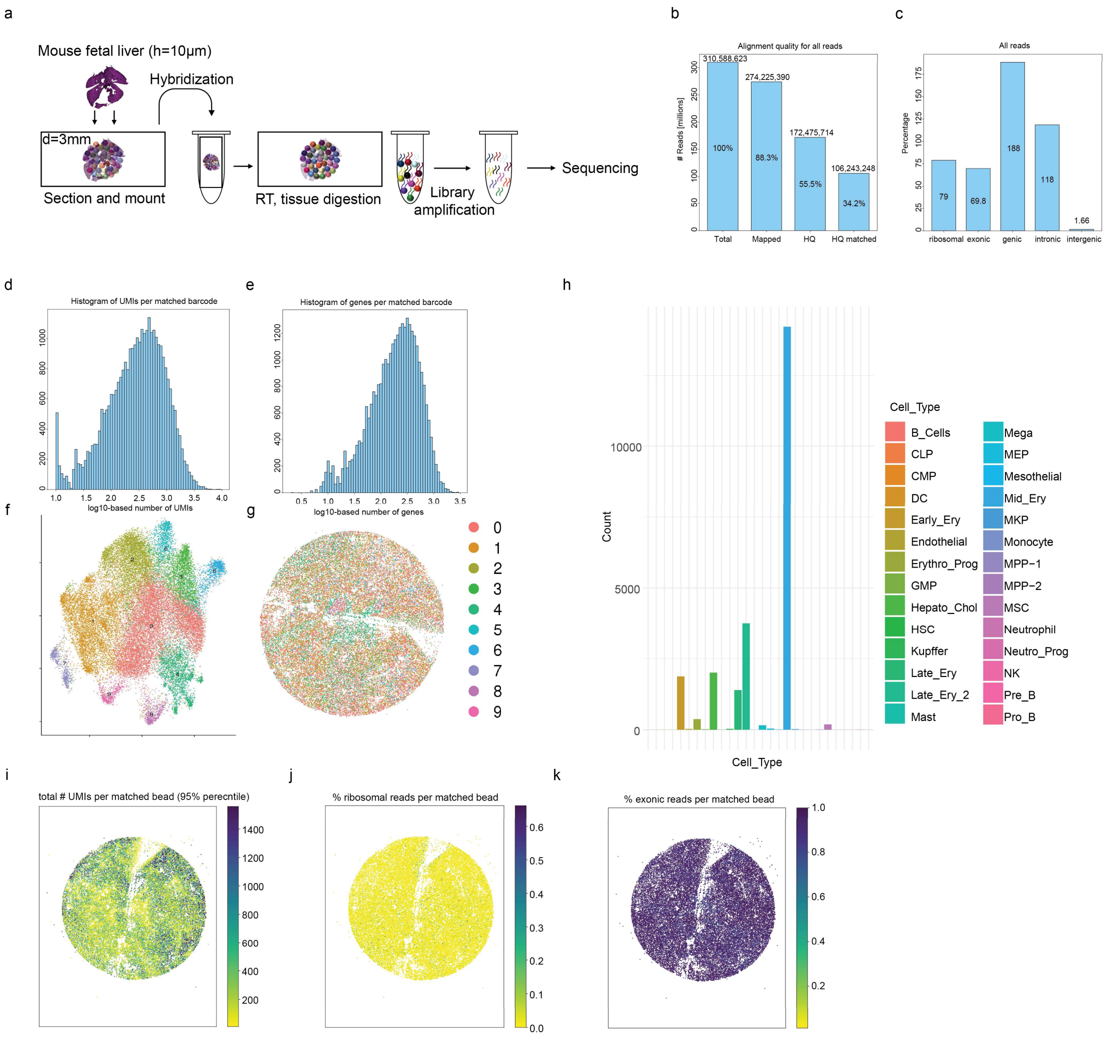


**Extended Data Figure 2. The workflow of Slide-seq on mouse fetal liver and the UMIs, genes, and reads from one of the E14.5 pucks**

(a) Illustrations depicting the workflow of Slide-seq.

(b) Bar graph showing the alignment quality for all reads.

(c) Bar graph showing the percentage of reads.

(d)(e) Histogram showing the UMIs (d) and genes (e) per matched.

(f) (g) UMAP (f) and Spatial UMAP (g) by RCTD clustering.

(h) Bar graph showing the count of signature mRNAs of each cell type identified.

(i)(j)(k) Spatial UMAP showing total UMIs(i), ribosomal reads(j), and exonic reads (k) per matched bead.

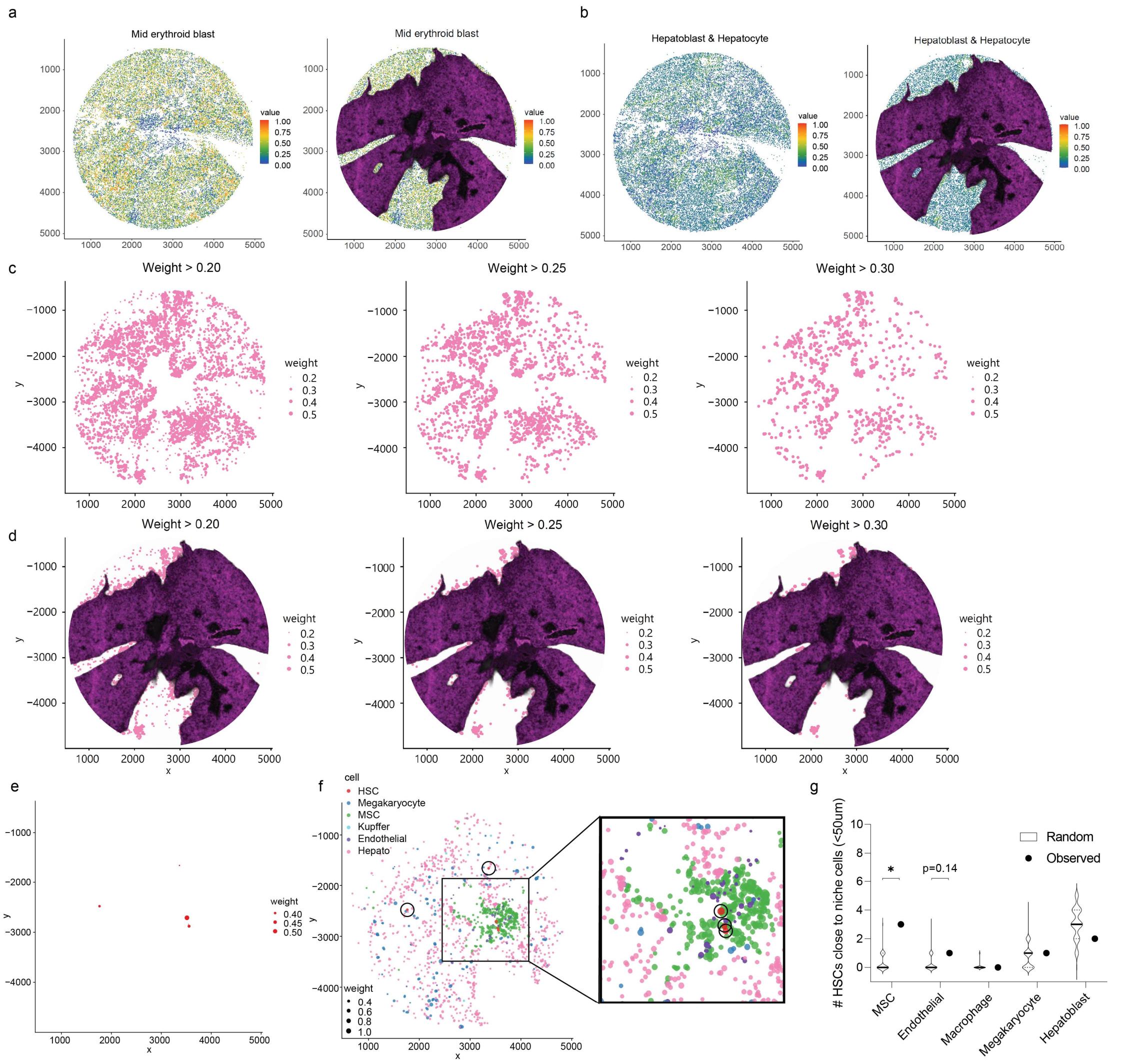


**Extended Data Figure 3. Mapping cellular architecture and interactions in E14.5 mouse fetal liver using Slide-seq.**

(a) Spatial UMAP showing the weight of signature genes for erythroid blasts on each bead.

(b) Spatial UMAP showing the weight of signature genes for hepatoblasts and hepatocytes on each bead.

(c) (d) Spatial UMAP showing the annotation of hepatoblasts and hepatocytes using different thresholds without (c) or with (d) tissue alignment. Aligned tissue was from a DAPI-stained adjacent pre-puck section.

(e) Identification of HSCs on another duplicated puck.

(f) Cellular architecture of E14.5 mouse fetal liver on another duplicated puck. Circles highlight the fetal liver HSCs.

(g) Permutation test demonstrating the enrichment of HSCs in the vicinity of potential niche cells corresponding to (f). The closest niche cell within 50μm to HSCs was considered as the niche cell to HSCs.

*, $p<0.05$. showing combined p-value calculated using Fisher’s combined probability test.

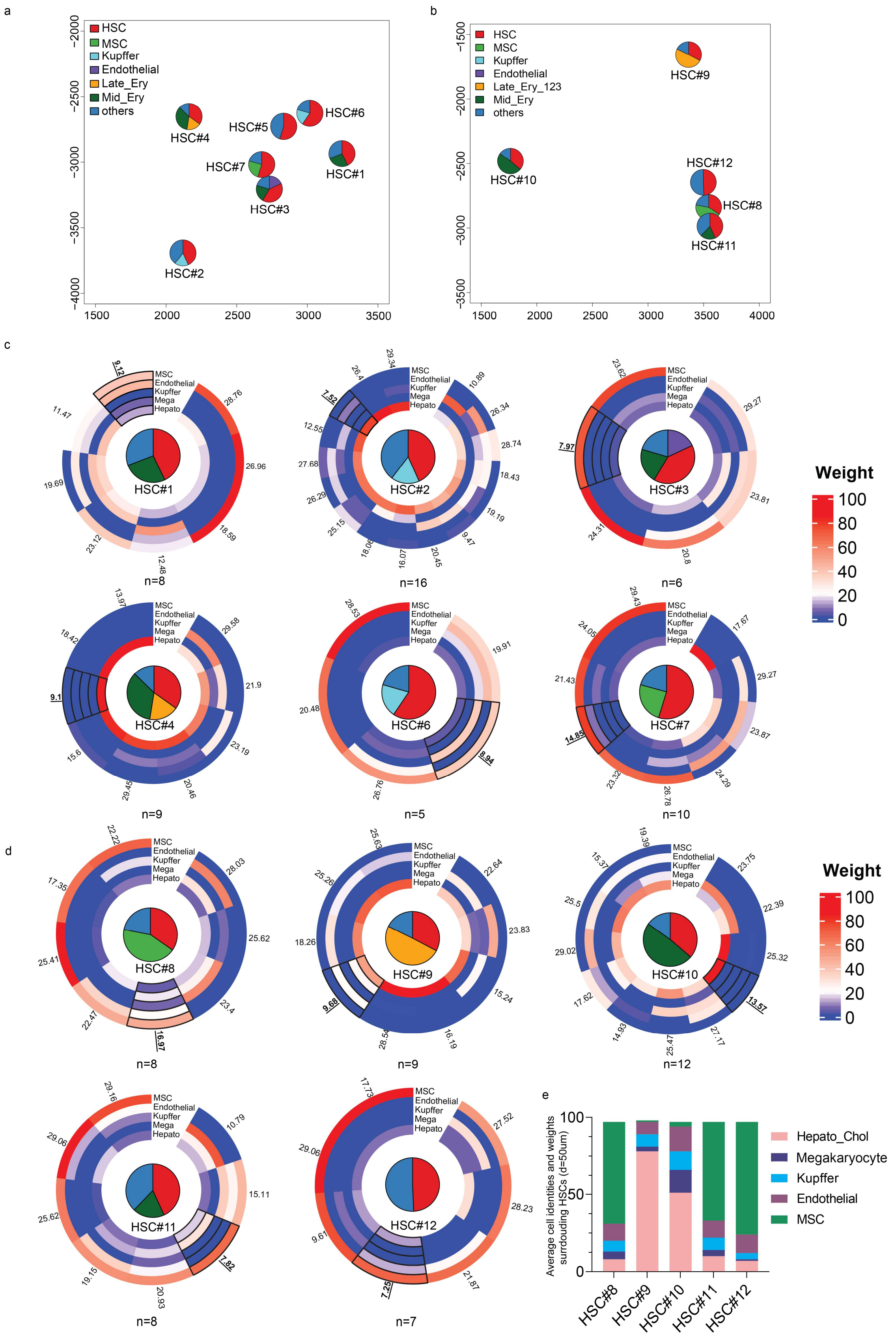
a
HSC
MSC
Kupffer
Endothelial
Late_Ery
Mid_Ery
others
HSC#1
HSC#2
HSC#3
HSC#4
HSC#5
HSC#6
HSC#7
−2000
−2500
−3000
−3500
−4000
1500
2000
2500
3000
3500
b
HSC
MSC
Kupffer
Endothelial
Late_Ery_123
Mid_Ery
others
HSC#8
HSC#9
HSC#10
HSC#11
HSC#12
−1500
−2000
−2500
−3000
−3500
1500
2000
2500
3000
3500
4000
c
MSC
Endothelial
Kupffer
Mega
Hepato
HSC#1
n=8
HSC#2
n=16
HSC#3
n=6
HSC#4
n=9
HSC#6
n=5
HSC#7
n=10
Weight
100
80
60
40
20
0
d
HSC#8
n=8
HSC#9
n=9
HSC#10
n=12
HSC#11
n=8
HSC#12
n=7
Weight
100
80
60
40
20
0
e
Average cell identities and weights surrouding HSCs (d=50um)
100
50
0
HSC#8
HSC#9
HSC#10
HSC#11
HSC#12
Hepato_Chol
Megakaryocyte
Kupffer
Endothelial
MSC

**Extended Data Figure 4. Decomposition analysis showing the overall niche cell identities surrounding each HSC**

(a) (b) Pie chart showing the weight of various sets of signature genes corresponding to different cell types on beads annotated as HSCs (HSC weight>35%)

(c) (d) Pie chart showing the overall cell identities and weights surrounding each HSC within 30μm. n = the number of beads within 30um of each HSC. For each figure, the inner pie chart shows the weight of various sets of signature genes corresponding to different cell types on beads annotated as HSCs (HSC weight>35%). Each of the outer concentric rings is representative of a specific niche cell identity and its weight on the corresponding bead. Numbers along the perimeter indicate the distance (μm) between the bead and the HSC. Bold borders were used to highlight the cell identifies and corresponding weights on the closest bead to each HSC. HSC #5 has only 1 bead (endothelial cells weight >99%) within 30μm. The weights of erythrocytes and erythroblasts were excluded on all the beads.

(e) Bar graph presenting the average cell identities within 50μm of each HSC from another duplicated puck. Decomposition was applied to all the beads within 50μm of each HSC. The weights of erythrocyte and erythroblasts were excluded on all the beads.

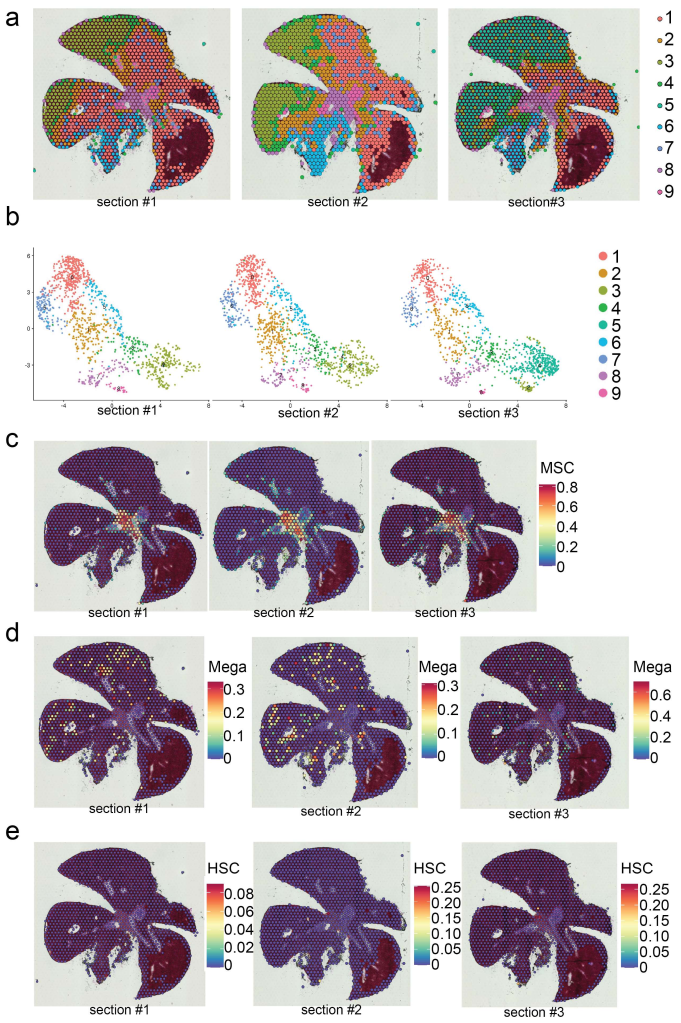


**Extended Data Figure 5. Identifying HSCs and niche cells in E14.5 mouse fetal liver using 10x Visium.**

(a) (b) Spatial UMAP (a) and UMAP (b) from three consecutive cryo-sections from E14.5 mouse fetal liver.

(c) (d) (e) Spatial UMAP identifying MSCs (c), Megakaryocytes (d), and HSCs (e) in 10x Visium after the application of RCTD.

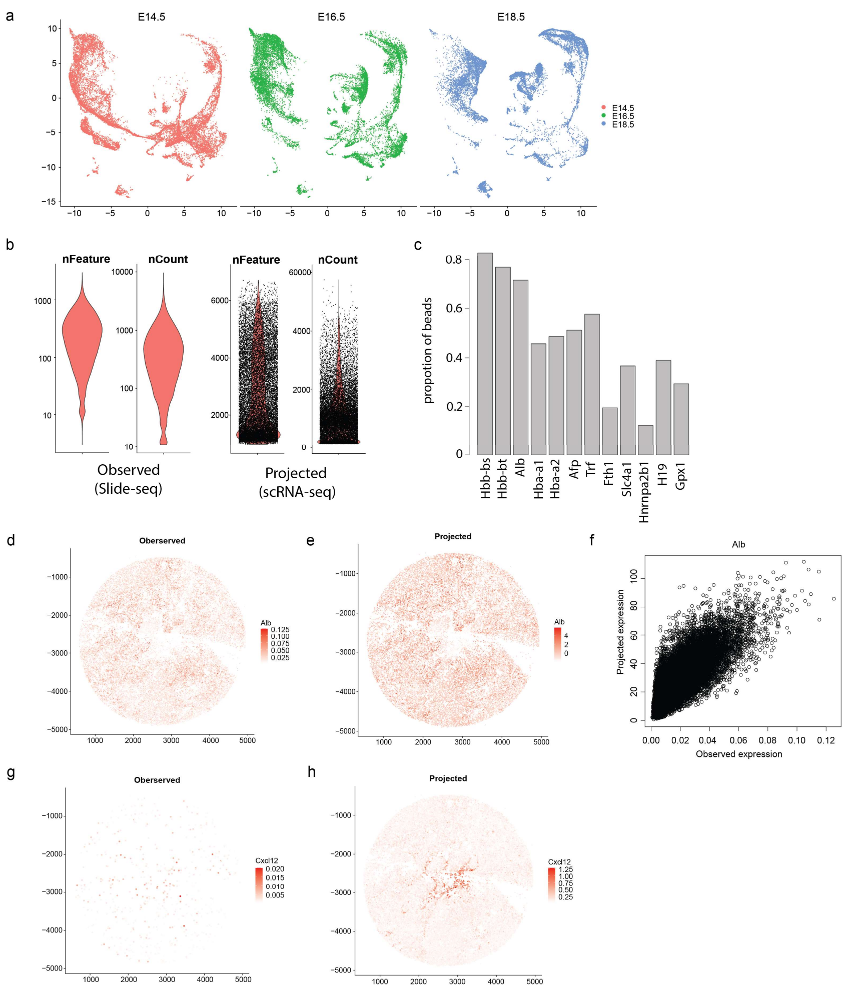


**Extended Data Figure 6. Enhancing Slide-seq gene expression using the projection method.**

(a) UMAP showing the cells from E14.5, E16.5, and E18.5 fetal liver.

(b) Violin plot showing the number of features and counts for Slide-seq and scRNA-seq.

(c) Bar graph showing the most abundant genes identified on the puck

(d) (e) Spatial UMAP showing the observed expression (d) and projected expression (e) of *Alb* on one of the pucks.

(f) The correlation between observed *Alb* expression (X-Axis) and projected *Alb* expression (Y-Axis).

(g) (H) Spatial UMAP showing observed *Cxcl12* expression (d) and projected *Cxcl12* expression (e) on one of the pucks.

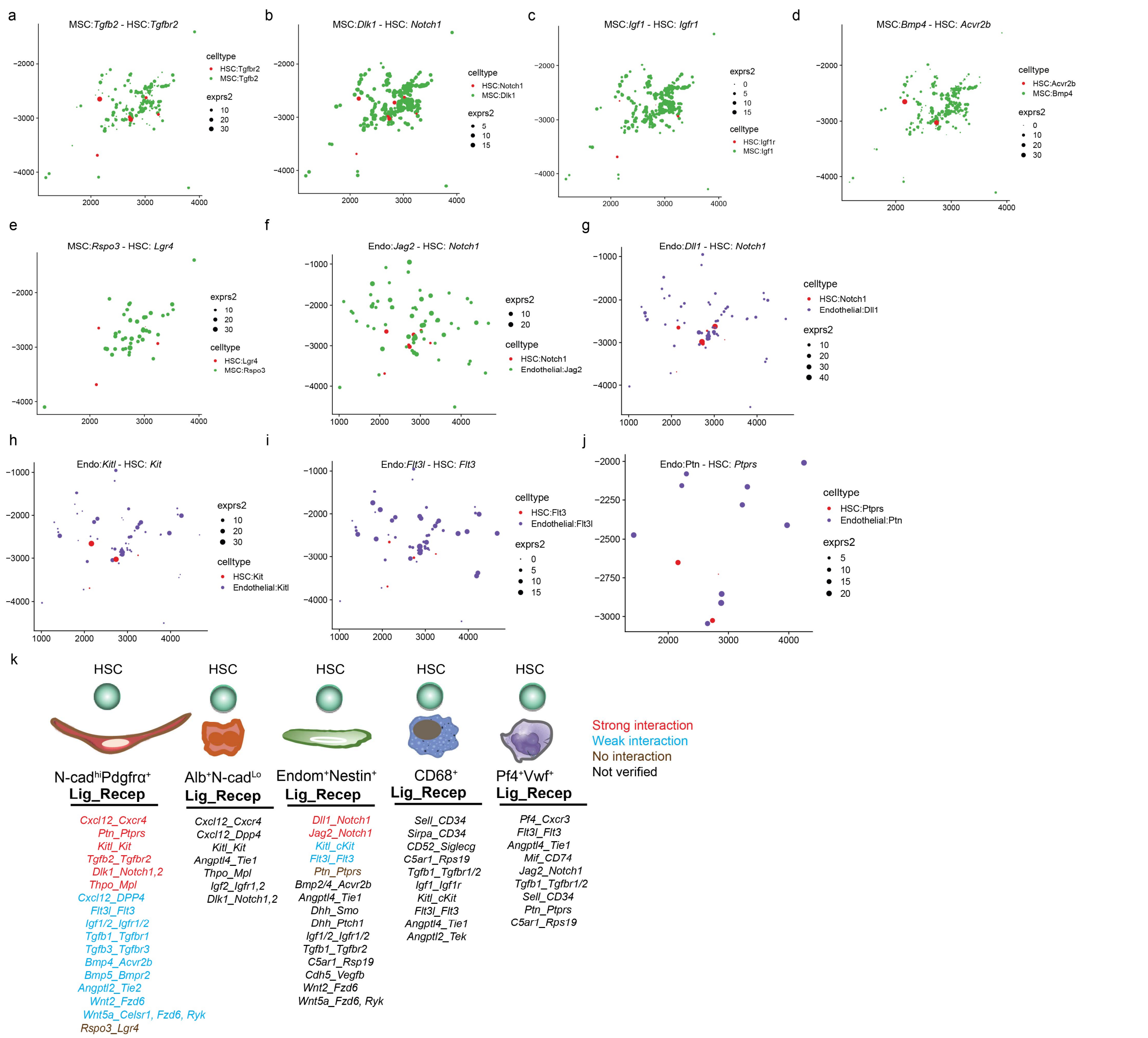


**Extended Data Figure 7. Validating and categorizing signaling between HSCs and the niche using Slide-seq**

(a) - (e) Spatial UMAP Showing the ligand-receptor interactions *Tgfb2-Tgfbr2* (a), *Dlk1-Notch1* (b), *Igf1-Igfr1* (c), *Bmp5-Bmpr2* (d), and *Rspo3-Lgr4* (e) between MSCs and HSCs.

(f) - (j) Spatial UMAP Showing the ligand-receptor interactions *Jag2-Notch1* (f), *Dll1-Notch1* (g), *Kitl-Kit* (h), *Fli3l-Flt* (i), and *Ptn-Ptprs* (j) between endothelial cells and HSCs.

(k) A summary of potential ligand-receptor interactions between niche cells and HSCs, predicted by CPDB. The interactions between MSCs and HSCs were further validated using Slide-seq on one of the pucks. Strong interactions were defined as those in which at least 3 out of 7 HSCs exhibits the ligand-receptor interaction with MSCs; Weak interactions were defined as those in which 1 or 2 out of 7 HSCs exhibits the ligand-receptor interaction with MSCs; No interactions were defined as those in which 0 out of 7 HSCs exhibits the ligand-receptor interaction with MSCs.

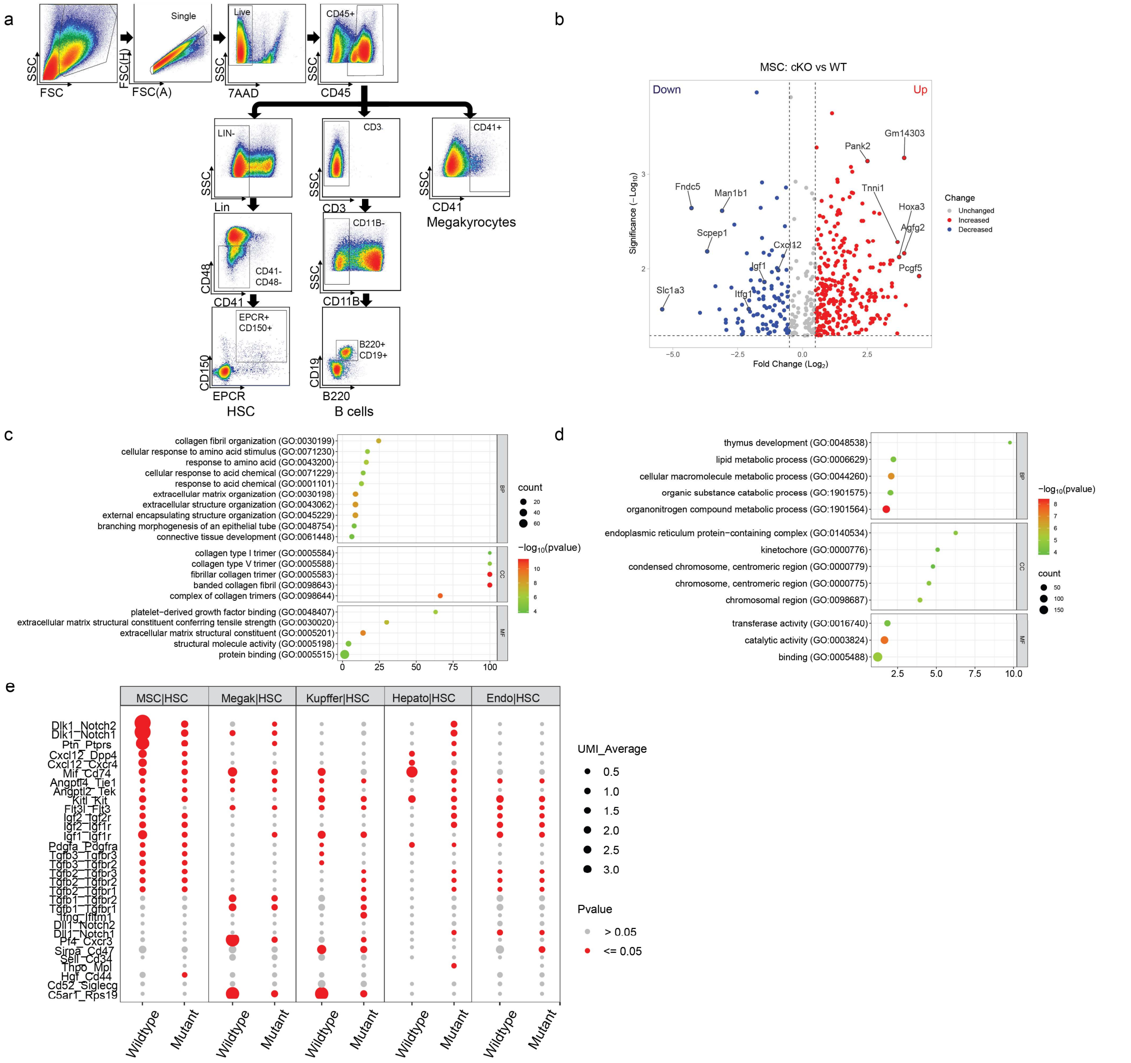


**Extended Data Figure 8. Conditional knockout of Cxcl12 from N-cad- expressing cells leads to the remodeling of MSCs.**

(a) The gating strategy for flow cytometry analysis and FASC.

(b) Volcano plot displaying differentially expressed genes (DEGs) in MSCs mediated by Cxcl12 cKO from N-cad-expressing cells.

(c) (d) Bubble plot illustrating the top GO terms in WT MSCs (c) and MSCs upon conditional knockout of Cxcl12 from N-cad-expressing cells (d) (BP: biological process; CC: cellular components; MF: Molecular Functions)

(e) CPDB analysis predicting the potential ligand-receptor changes in response to conditional knockout of from N-cad-expressing cells.

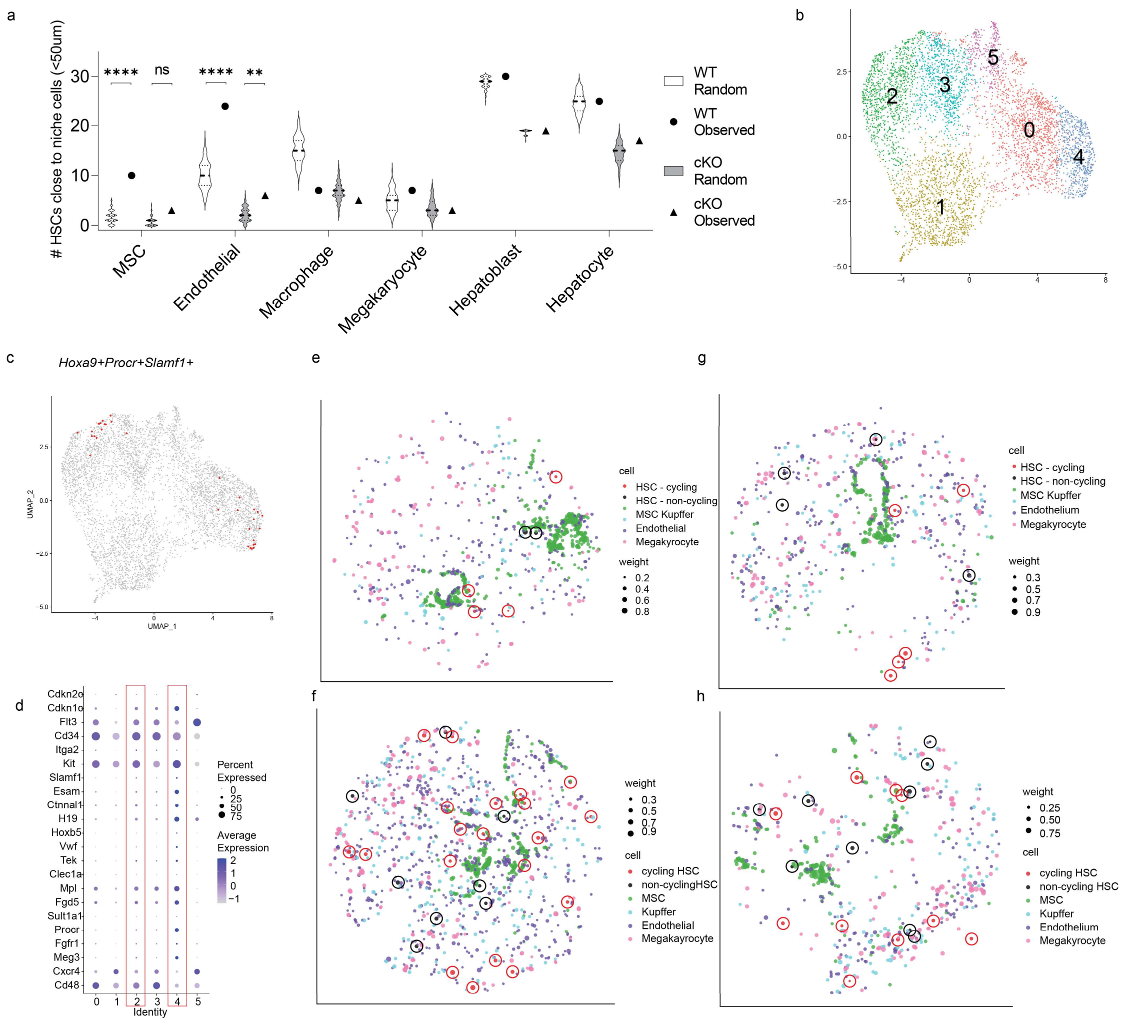


**Extended Data Figure 9. Conditional knockout of Cxcl12 from N-cad-expressing cells leads to the relocation of both cycling HSCs and non-cycling HSCs**

(a) Permutation test demonstrating total HSC enrichment in the vicinity potential niche cells in WT and cKO mice on another duplicated puck.

(b) Sub-clustering of HSCs in E16.5 scRNA-seq

(c) UMAP showing *Hoxa9*$^{+}$*Slamf1*$^{+}$*Procr*$^{+}$ HSCs in (b).

(d) DEG analysis showing the signature genes in cycling HSCs (cluster 2) and non-cycling HSCs (cluster 4).

(e) - (h) Spatial UMAP displaying the cellular architecture in E16.5 mouse fetal liver from wild-type (WT) (e) (f) and conditional knockout (cKO) (g) (h) mice (hepatoblasts and hepatocytes not shown due to the high abundancy). The red circles highlight

cycling HSCs; the black circles highlight non-cycling HSCs.
**, $p<0.01$; ****, $p<0.0001$. showing combined p-value calculated using Fisher's combined probability test.

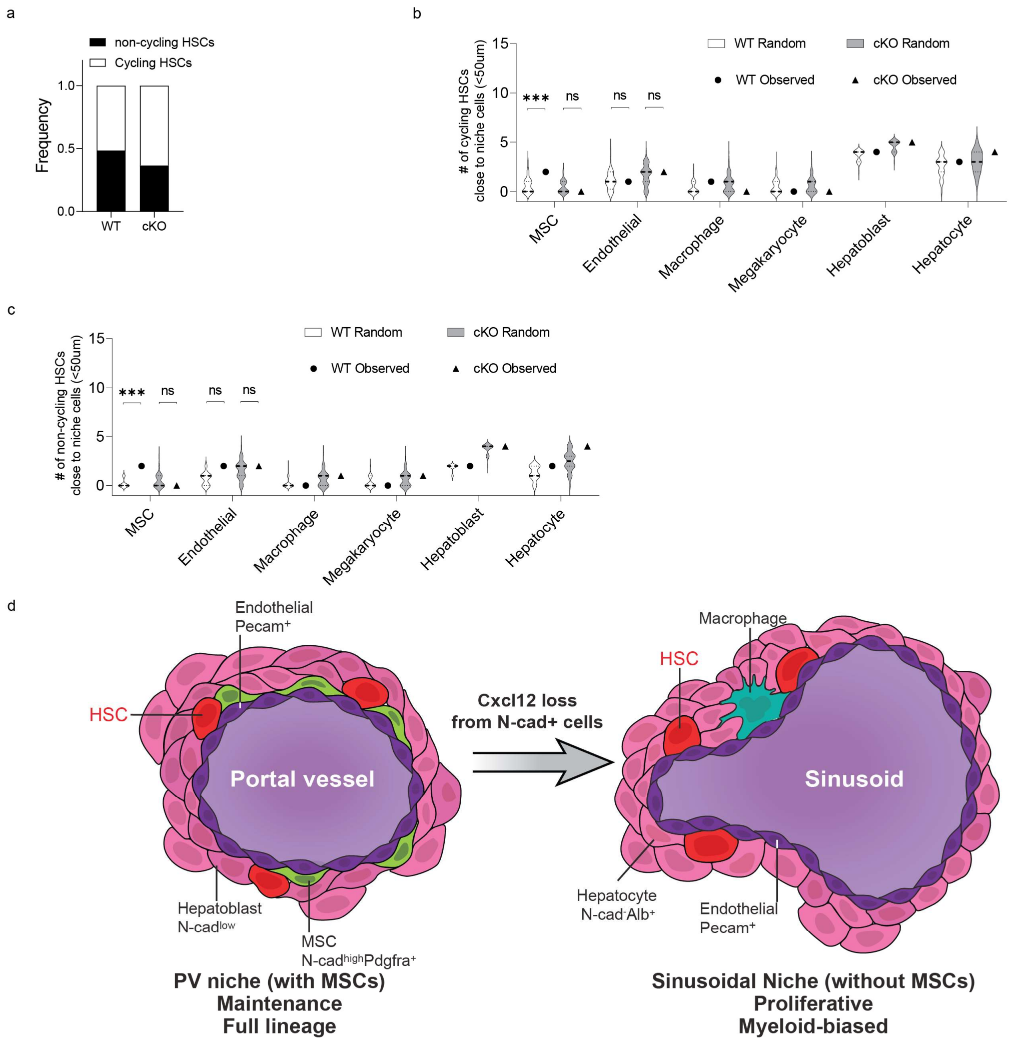


**Extended Data Figure 10. Postulated mechanism of the cellular structure of distinct niches and their role in supporting FL-HCSs.**

(a) Bar plot illustrating the proportions of cycling HSCs, and non-cycling HSCs in WT and cKO mice.

(b) (c) Permutation test demonstrating the enrichment of cycling HSCs (b), and non-cycling HSCs (c) in proximity to potential niche cells in WT and cKO mice.

(d) Postulated mechanism of the cellular structure of distinct niches and their role in supporting FL-HCSs.

***, p<0.001. showing combined p-value calculated using Fisher's combined probability test.

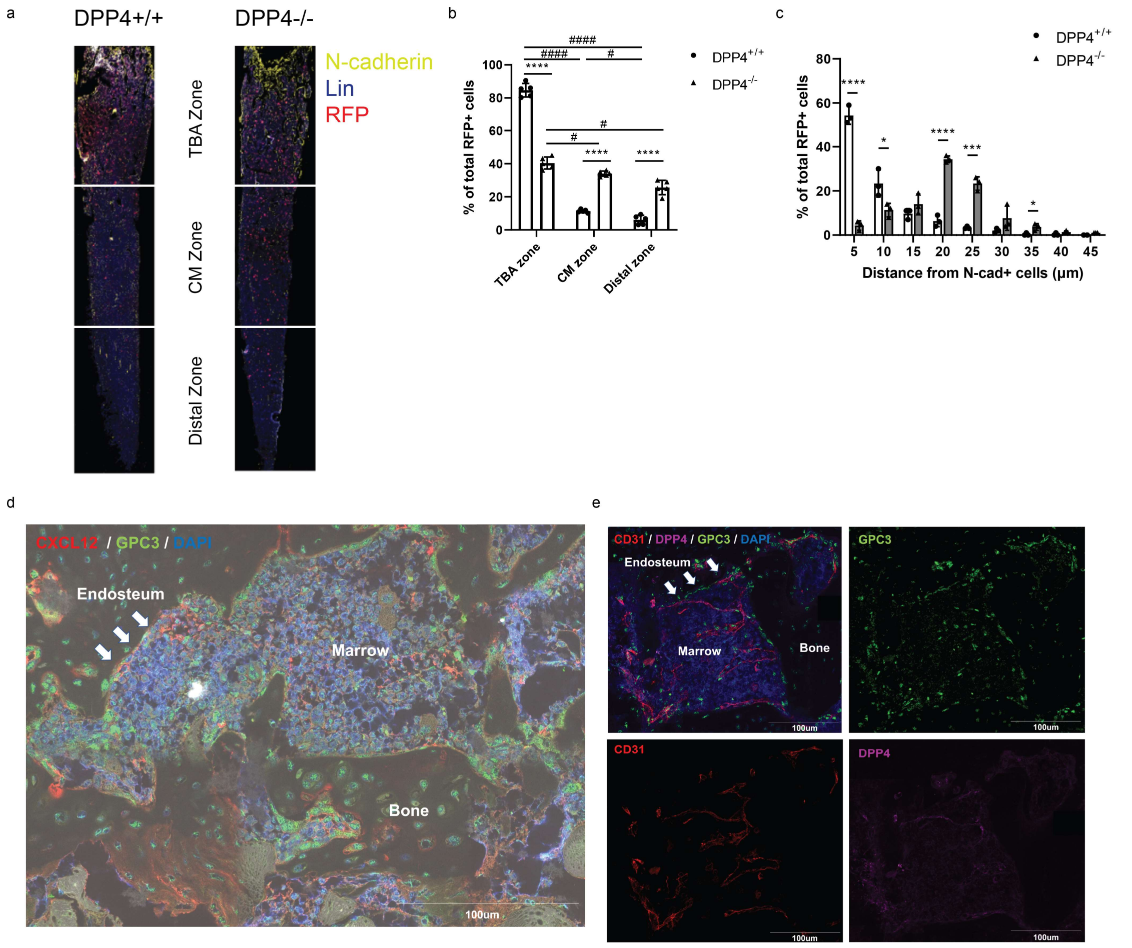


**Extended Data Figure 11. Gpc3 and Dpp4 potentially modulate the HSPCs homing to the trabecular bone area in adult mouse.**

(a) Immunofluorescent staining of RPF (red), lineage (blue), and N-cadherin (yellow) showing the distribution of donor bone marrow cells (from Dpp4$^{+/+}$ or Dpp4$^{-/-}$ mT/mG mice) in the recipient bone marrow after 4 weeks.

(b) Bar plot indicating the distribution of RPF+ donor bone marrow cells (from Dpp4$^{+/+}$ or Dpp4$^{-/-}$ mT/mG mice) in the three different zones of recipient bone marrow.

(c) Bar plot showing the distance of RPF+ donor bone marrow cells (from Dpp4$^{+/+}$ or Dpp4$^{-/-}$ mT/mG mice) to N-cadherin+ cells in mouse bone marrow

(d) Immunofluorescent staining of Cxcl12 (red), Gpc3 (green), and DAPI (blue) in the TBA of WT mouse bone marrow.

(e) Immunofluorescent staining of Cd31 (red), Dpp4(purple), Gpc3 (green), and DAPI (blue) in the TBA of WT mouse bone marrow.

*, $p<0.05$; ***, $p<0.001$; ****, $p<0.0001$ showing p-value calculated using student t's test.

####,$p<0.0001$ showing p-value calculated using one-way ANOVA test.
